# Non-autonomous regulation of germline stem cell proliferation by somatic MPK-1/MAPK activity in *C. elegans*

**DOI:** 10.1101/2020.08.24.265249

**Authors:** Sarah Robinson-Thiewes, Benjamin Dufour, Pier-Olivier Martel, Xavier Lechasseur, Amani Ange Danielle Brou, Vincent Roy, Yunqing Chen, Judith Kimble, Patrick Narbonne

## Abstract

Extracellular signal-regulated kinase (ERK)/mitogen-activated protein kinase (MAPK) is a major positive regulator of cell proliferation that is often upregulated in cancer. Yet few studies have addressed ERK/MAPK regulation of proliferation within a complete organism. The *C. elegans* ERK/MAPK ortholog MPK-1 is best known for its control of somatic organogenesis and germline differentiation, but it also stimulates germline stem cell proliferation. Here we identify tissue-specific MPK-1 isoforms and characterize their distinct roles in germline function. The germline-specific MPK-1B isoform promotes germline differentiation, but has no apparent role in germline stem cell proliferation. By contrast, the soma-specific MPK-1A isoform promotes germline proliferation non-autonomously. Indeed, MPK-1A functions in the intestine or somatic gonad to promote germline proliferation, independently of its other known roles. We propose that a non-autonomous role of ERK/MAPK in stem cell proliferation may be conserved across species and other tissue types, with major clinical implications for cancer and other diseases.

## Introduction

The extracellular signal-regulated kinase (ERK)/mitogen-activated protein kinase (MAPK) is the downstream effector of a conserved oncogenic pathway that is amongst the most often upregulated in cancer^1-3^. Conversely, mutations that inactivate ERK/MAPK are seldom found in tumours, emphasizing the key role of this kinase in proliferation^2, 4^ The prevailing paradigm is that activated ERK/MAPK functions cell autonomously, responding to growth factors in the same cells where it promotes proliferation^5, 6^. However, the role of ERK/MAPK in stem cells, including mammalian embryonic stem cells, has been perplexing. Most evidence indicates that ERK/MAPK is dispensable for stem cell proliferation, but is required for differentiation^7-10^. Although loss of ERK1/2 in mammalian embryonic stem cells reduces their proliferation in culture, that effect may be secondary to genomic instability^11-13^. The effect of ERK/MAPK on stem cell proliferation in an organism remains largely unexplored.

We have investigated how ERK/MAPK affects stem cell proliferation in *C. elegans*. The genome of this small nematode possesses a single ortholog of mammalian ERK1/2 MAPKs, MPK-1, which has been investigated for years by classical genetic analyses. In *mpk-1(ø)* null mutants, several somatic organs fail to develop normally (*e.g*. vulva)^14-16^, and the germline fails to progress through the pachytene stage of the meiotic cell cycle, which causes sterility^17-19^. The role of MPK-1 in germline proliferation, however, has been puzzling. Proliferation is reduced in *mpk-1(ø)* mutants^18, 20, 21^, but also reduced upon removal of LIP-1, an MPK-1 inhibitory phosphatase that is expressed in germline stem cells (GSCs)^22^. Regardless, germline proliferation continues into adulthood in *mpk-1(ø)* mutants^21^. Therefore, as in other species and other stem cell types, nematode ERK/MAPK is essential for differentiation but not for stem cell proliferation.

The *C. elegans* adult gonad has a simple architecture. Two U-shaped gonadal arms house the germline (Fig. 1A, left), where germ cell maturation occurs along a distal-proximal axis (Fig. 1A, right). A pool of proliferative GSCs is maintained at the distal end within a somatic niche; as GSC daughters leave the niche, likely displaced by proliferation, they begin differentiation and progressively mature into gametes at the proximal end^20, 23-26^. GSCs, together with their proliferating progeny, span a region in the distal gonad termed the progenitor zone (PZ) (Figure 1A, right). The rate of GSC proliferation is inferred from the mitotic index measured across the PZ^20, 21, 27, 28^. In adult hermaphrodites, GSC proliferation rates influence distal-to-proximal germ cell flow, and are thus linked to the pace of oocyte production^29^. Once germ cells leave the progenitor zone, they proceed through meiotic prophase and undergo gametogenesis, making sperm as larvae and oocytes in adults.

**Figure 1.**
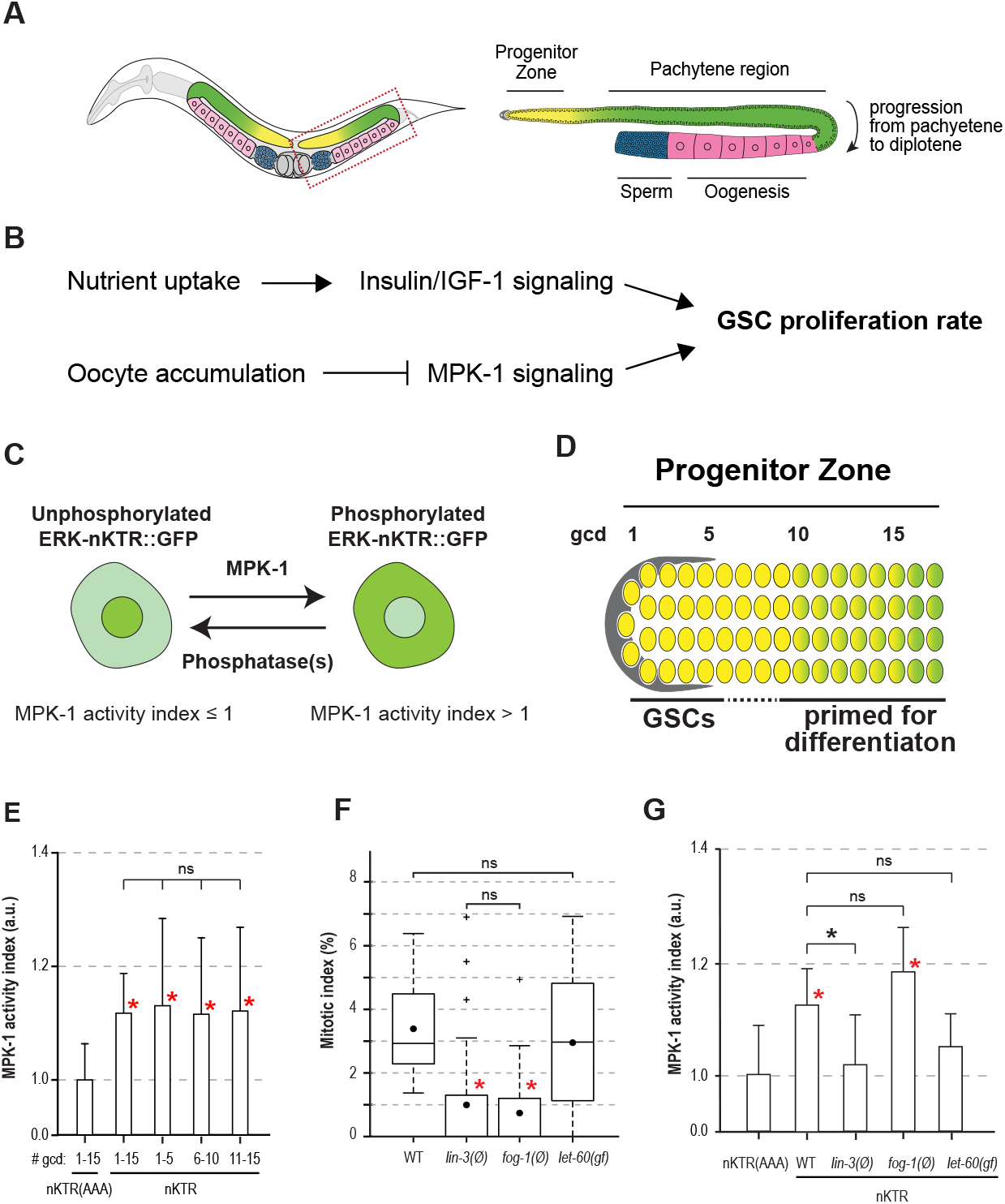
GSC MPK-1 activity does not correlate with GSC proliferation. (**A**) Left, schematic of an adult *C. elegans* hermaphrodite with posterior gonadal arm boxed; right, gonadal arm labelled with regions relevant to this work. (**B**) GSC proliferation rate is regulated by both Insulin/IGF-1 signaling and MPK-1 signaling, which act in parallel with additive effects^21^. (**C**) Schematic of *in vivo* assay for MPK-1 activity. Sensor GFP is enriched in the nucleus when MPK-1 activity is absent (left), but becomes cytoplasmic upon phosphorylation by MPK-1/ERK (right). The MPK-1 index refers to the ratio of cytoplasmic to nuclear GFP, normalized to the ERK-nKTR(AAA) baseline control. An index >1 indicates MPK-1 kinase activity. (**D**) Proliferating germ cells in the Progenitor Zone include a pool of GSCs within the niche (grey) and GSC daughters that have launched the differentiation program but not yet begun overt differentiation. Numbers mark positions along the germline axis in germ cell diameters (gcd) from the distal end. (**E**) MPK-1 activity in the Progenitor Zone. ERK-nKTR is cytoplasmically-enriched compared to the ERK-nKTR(AAA) baseline control. The MPK-1 index is similar across the PZ. Red asterisk indicates statistical significance of ERK-nKTR at each binned position compared to baseline ERK-nKTR(AAA) control (p < 0.01); determined by ANOVA followed by Tukey; ns, not significant. Error bars indicate standard deviation. Sample sizes are 25 animals each for ERK-nKTR and ERK-nKTR(AAA), scoring 5 PZ cells per animal (see methods). (**F**) Box plots of mitotic indices in wild-type (WT) and mutant Progenitor Zones. Dots mark averages. Red asterisk indicates statistical significance compared to WT (p < 0.01; Kruskal-Wallis followed by Dunn). Sample sizes, from left to right, are 14, 39, 17, 19. (**G**) MPK-1 activity is lost from the Progenitor Zone in *lin-3(ø)* and *let-60(gf)*, but not *fog-1(ø)* mutants. For simplicity, ERK-nKTR(AAA) data are shown only for WT (see Fig. S1 for all raw ratios). Error bars indicate standard deviation. Red asterisks as in **E**; a black asterisk indicates statistical significance in pairwise comparisons (p < 0.01). Sample sizes, for each (ERK-nKTR(AAA); ERK-nKTR) pair, are WT (25, 25), *lin-3(ø)* (7, 10), *fog-1(ø)* (11, 12), *let-60gf* (10, 10), scoring 5 PZ cells per animal.

*C. elegans* GSC proliferation rates are controlled jointly by ERK/MAPK (see above) and insulin/IGF-1 signaling (IIS) (Fig. 1B). IIS promotes stem cell proliferation downstream of nutrient uptake (Fig. 1B), a regulation conserved in *C. elegans, Drosophila* and likely mammals^30-36^. Most relevant here, downstream IIS effectors act autonomously within the germline^21, 34^. In parallel, but through an unknown mechanism, MPK-1 signaling combines with IIS to promote the high proliferation typical of young adult hermaphrodites (Fig. 1B)^21^. One aspect of MPK-1’s role in germline proliferation was clarified through studies of homeostatic regulation. That is, GSC proliferation plummets in well-fed hermaphrodites that accumulate unfertilized oocytes because of a lack of sperm^21, 27, 37, 38^. This homeostatic lowering of GSC proliferation occurs even though IIS remains systemically active, but requires inhibition of MPK-1 signaling^21, 27^ This control occurs specifically in the sperm-depleted gonad arm^27^ and therefore must be localized to that arm. Importantly, MPK-1 signaling inhibition is central to homeostatic regulation but the cells or tissues involved have not been defined. Although molecular details may differ, homeostatic regulation is a common phenomenon found, for example, in *Drosophila* hematopoietic and gut stem cells, and mammalian hair follicle stem cells^39-41^.

An outstanding question remains how ERK/MAPK controls stem cell proliferation. Our primary focus in this work is to understand how *C. elegans* MPK-1 promotes germline stem cell proliferation. We find that MPK-1 acts in the animal’s gut and/or somatic gonad to regulate the rate of GSC proliferation and that its action is non-autonomous. This finding challenges the prevailing view that ERK/MAPK acts autonomously to control proliferation. It therefore has potential to impact our understanding of ERK/MAPK controls during development, homeostatic stem cell regulation, and tumour formation with implications for all organisms.

## Results

### MPK-1 has low but significant kinase activity in wild-type GSCs

MPK-1 promotes the high rate of GSC proliferation typical of young adult hermaphrodites^21^. While MPK-1 protein is present in GSCs, its catalytically active form, diphosphorylated MPK-1 (dpMPK-1), has only been detected in meiotic germs cells and oocytes^18, 19, 42^. This apparent lack of active MPK-1 in GSCs led us to investigate how MPK-1 affects germline proliferation. We considered three hypotheses: MPK-1 has a kinase-independent role in germ cell proliferation; active MPK-1 is present in GSCs but only at a previously undetected level; or MPK-1 promotes proliferation non-autonomously, from outside GSCs. To distinguish among these possibilities, we first used a sensitive *in vivo* ERK nuclear kinase translocation reporter (ERK-nKTR) to reassess MPK-1 activity in GSCs^43^. This GFP-tagged sensor harbors three MPK-1-specific phosphorylation sites that control its subcellular localization. When the reporter is unphoshorylated, GFP is retained within the nucleus, but when phosphorylated, GFP is exported to the cytoplasm (Fig. 1C). The ratio of cytoplasmic to nuclear GFP therefore provides an index of MPK-1 activity. A control reporter that lacks MPK-1 sites, ERK-nKTR(AAA), was used to establish the baseline ratio of cytoplasmic to nuclear GFP in each experiment^43^. Ratios above this baseline were scored as the MPK-1 activity index (see Methods for details).

We first assessed MPK-1 activity in the Progenitor Zone of wild-type adult hermaphrodites. Specifically, MPK-1 activity index was determined in PZ cells grouped by distance from the distal end (Fig. 1D). Indices were higher than baseline and similar throughout the PZ (Figs. 1E-F, S1). Our assay also detected the well-established activity in pachytene germ cells and oocytes (Fig. S1)^18, 19, 42, 43^. The discovery of active MPK-1 in GSCs raised the possibility that MPK-1 acts autonomously to promote GSC proliferation.

We took advantage of three mutants to ask whether MPK-1 activity in GSCs is correlated with their proliferation. Such a correlation would be consistent with MPK-1 autonomous control of germline proliferation. The first two mutants, *lin-3(ø)* and *fog-1(ø)*, have a lowered rate of GSC proliferation due to homeostatic feedback: *lin-3(ø)* mutants accumulate endomitotic oocytes because they cannot ovulate, while *fog-1(ø)* mutants accumulate unfertilized oocytes because they cannot make sperm (Fig. S2)^27, 44, 45^. As expected, the mitotic index was very low in both mutants (Fig. 1F). MPK-1 activity in the PZ, however, was not the same in the two mutants. It was undetectable in *lin-3(ø)*, but similar to wild-type in *fog-1(ø)* mutants (Fig. 1G). Therefore, the presence of MPK-1 activity in the PZ can be uncoupled from GSC proliferation. We next examined a mutant predicted to have higher than normal MPK-1 activity, a gain-of-function *(gf)* mutant in *let-60*, which encodes Ras, a central MPK-1 activator^16, 17^ Surprisingly, MPK-1 activity in the PZ was undetectable in *let-60(gf)* (Fig. 1G). Yet GSC proliferation in *let-60(gf)* was about the same as wild-type (Fig. 1F). This result also uncouples MPK-1 activity in the PZ from GSC proliferation. We conclude therefore that the level of MPK-1 activity in GSCs appears to be unrelated to their proliferation and therefore unlikely to be causative.

We next considered the possibility that MPK-1 might promote GSC proliferation from a different region within the germline tissue. This idea emerges from the fact that MPK-1 activity is stronger in the pachytene and oocyte regions than in the PZ. Indeed, MPK-1 activity levels in pachytene and oocytes of the selected mutants was correlated with their GSC MIs (Figs. 1F, S1L-M). Therefore, a non-autonomous action of MPK-1 from one germline region to another remains possible (though see below). Regardless, we conclude that MPK-1 activity in the GSCs themselves does not correlate with GSC proliferation

### Germline-specific MPK-1B isoform autonomously promotes fertility

To investigate how MPK-1 promotes GSC proliferation in more depth, we delineated both the expression and function of its two isoforms. The *mpk-1* locus encodes two transcripts: *mpk-1b* is the longer of the two and possesses a unique first exon, while *mpk-1a* shares all other exons with *mpk-1b* (Fig. 2A)^15, 16, 19^. Previous work demonstrated that *mpk-1b* is the main germline isoform, but did not fully address isoform-specific expression and germline function^19^. We first inserted epitope tags into the endogenous *mpk-1* locus using CRISPR/Cas9 gene editing^46^. A single V5 tag was inserted into the *mpk-1b*-specific exon to label MPK-1B protein specifically, and two tandem OLLAS tags were inserted into the shared C-terminus to label both MPK-1A and MPK-1B (Fig. 2A). The wild-type version of this dual tagged locus is called *mpk-1(DT)*. We next deleted 125 bp of the *mpk-1b*-specific exon in *mpk-1(DT)* to create an isoform-specific frameshift (henceforth called *mpk-1b(Δ)*) (Fig. 2A). In parallel, we created a 2221 bp in-frame deletion in *mpk-1(DT)* to remove common coding regions of the two isoforms (henceforth *mpk-1(Δ)*) (Fig. 2A). These alleles allowed us to unambiguously determine where each isoform is expressed and to define their respective biological roles.

**Figure 2.**
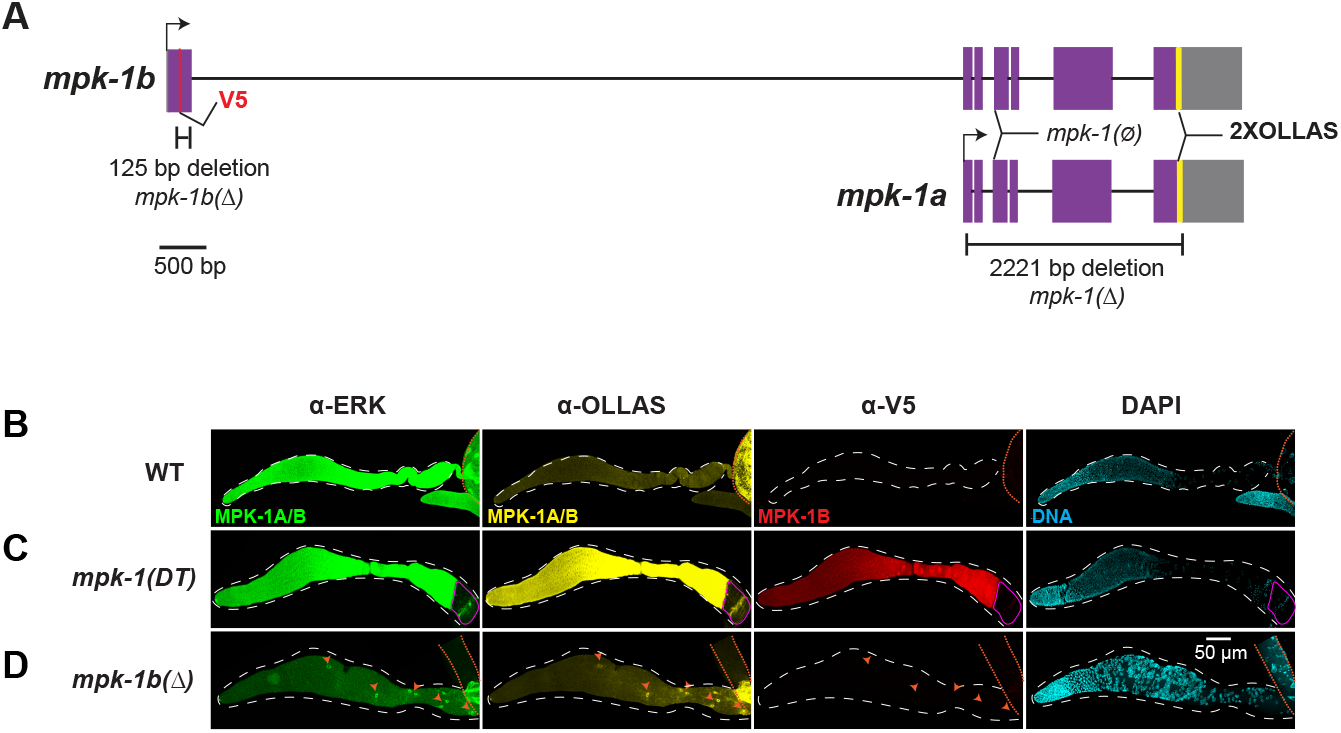
MPK-1B is germline-specific. (**A**) The two *mpk-1* isoforms. Purple boxes, coding exons; gray boxes, UTRs; lines connecting exons, introns; red line, V5 tag; yellow line, 2xOLLAS tags. The *mpk-1b* isoform possesses a unique first exon and a long first intron; *mpk-1a* and *mpk-1b* share all other exons, introns and the 3’UTR. Dual tagged *mpk-1(DT)* harbors a V5 tag in the *mpk-1b* specific exon that marks MPK-1B protein specifically and C-terminal 2xOLLAS tags that mark both MPK-1A and MPK-1B proteins. The *mpk-1b(Δ*) deletion removes most of the *mpk-1* specific exon and shifts the reading frame to eliminate MPK-1B. The *mpk-1(Δ*) deletion removes most of the shared exons and introns to eliminate MPK-1A and MPK-1B (see Methods). The *mpk-1(ga117)* is considered a null^18^ and shown here as *mpk-1(ø)*. (**B-D**) Dissected and stained adult gonads. When a stain does not distinguish between MPK-1A and MPK-1B, it is noted as MPK-1A/B. Anti-OLLAS staining, yellow; anti-V5 staining, red; anti-ERK staining, green; DAPI staining, cyan. White dashed line, boundaries of germline tissue; orange dashed line, boundaries of somatic tissue. Orange arrowheads, gonadal sheath nuclei. Pink outline, zygote.

We investigated expression of MPK-1A and MPK-1B in dissected hermaphrodite gonads that were stained with α-ERK, α-V5 and α-OLLAS antibodies plus DAPI. These gonad arms include the entire germline tissue plus several somatic gonadal cells, most prominently 10 sheath cells. Some preparations included the extruded gut in addition to the gonad. We used α-ERK to recognize MPK-1 on its own, with or without an epitope tag^18, 19^, and α-V5 and α-OLLAS to recognize epitope-tagged MPK-1. Wild-type gonads stained robustly with α-ERK, as previously reported^18, 19^, but not with α-V5 or α-OLLAS (Figs. 2B, S3A). The *mpk-1(DT)* gonads, on the other hand, stained robustly with all three antibodies (Figs. 2C, S3B-C). The intense germline staining in both wild-type and *mpk-1(DT)* precluded visualization of staining in somatic gonadal cells. However, the *mpk-1(DT)* gut showed strong α-ERK and α-OLLAS staining, but lacked α-V5 staining (Fig. S3C). We next stained *mpk-1b(Δ*) gonads, which do not have MPK-1B or the V5 tag, but do still have OLLAS-tagged MPK-1A. The intense α-ERK, α-OLLAS, and α-V5 signals were lost across the germline in *mpk-1b(Δ*) gonads (Fig. 2D). Moreover, α-ERK and α-OLLAS staining became visible in the somatic sheath cells (Figure 2D, orange arrowheads). Therefore, MPK-1A is the somatic isoform (expressed in gut and somatic gonad, but not germline), while MPK-1B is the germline isoform (expressed in germline, but not gut).

To investigate the biological function of the two isoforms, we first scored fertility and vulva formation. Wild-type and *mpk-1(DT)* animals were fertile, made a normal vulva and were otherwise indistinguishable (Fig. 3A). Thus, the tags have no apparent effect on MPK-1 function. By contrast, *mpk-1(Δ)* and *mpk-1b(Δ)* mutants were sterile and *mpk-1(Δ)*, but not *mpk-1b(Δ)*, mutants were vulvaless (Fig. 3A). The *mpk-1(Δ)* defects therefore match those of the established *mpk-1(ø)* null allele, *mpk-1(ga117)^15^*, but *mpk-1b(Δ)* do not. Together, these data indicate that the germline-specific MPK-1B functions autonomously to promote fertility since its removal results in sterility. Moreover, the well-formed vulva in *mpk-1b(Δ)* mutants indicates that somatic MPK-1A is responsible for vulva development, where it also acts autonomously in the vulva precursor cells^16^.

**Figure 3.**
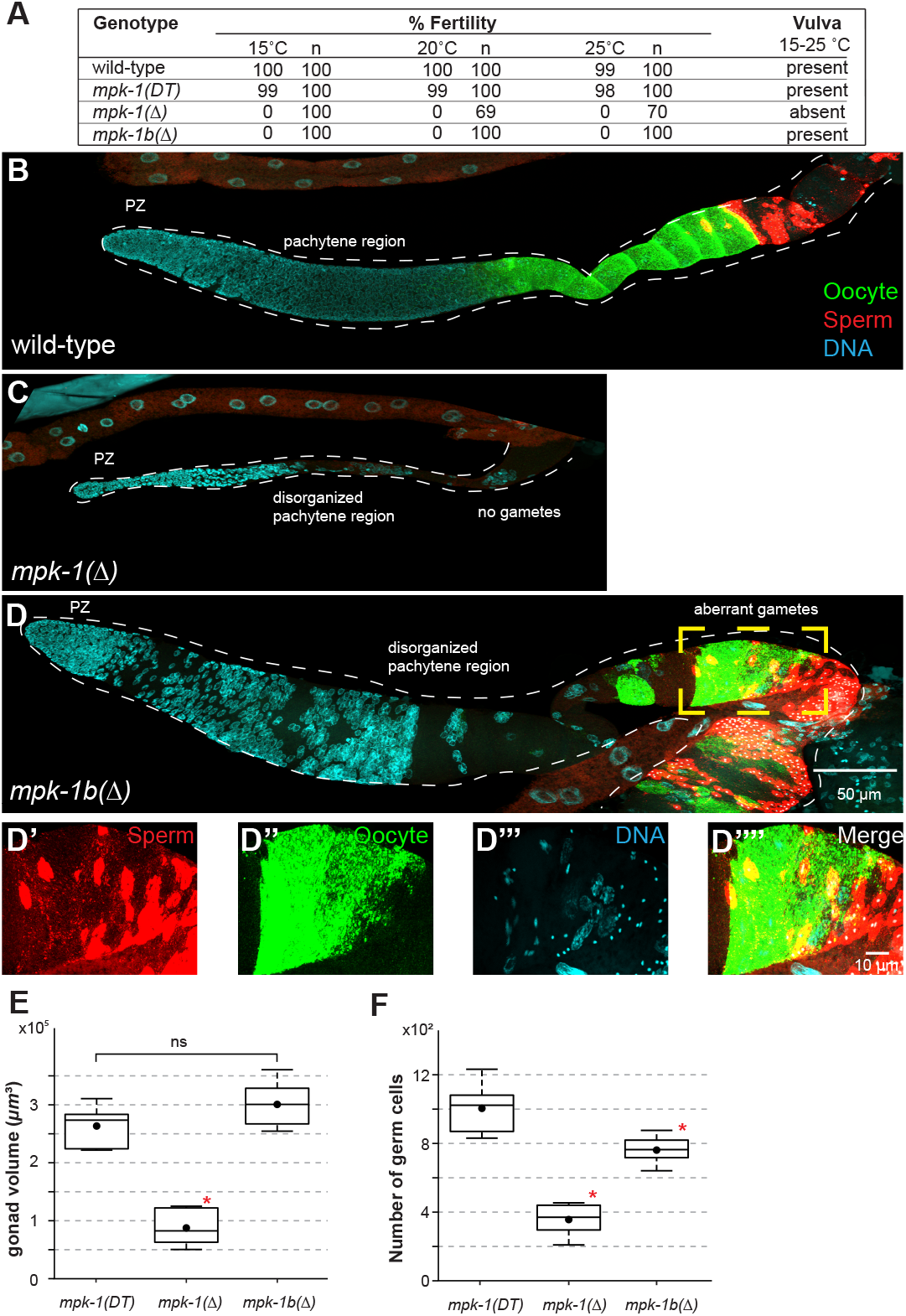
MPK-1B is required for fertility. (**A**) Wild-type and *mpk-1(DT)* are both fertile, while *mpk-1b(Δ)* and *mpk-1(Δ*) are both sterile. Briefly, all strains were maintained for at least 1 generation at each temperature before scoring for embryos (see Methods). (**B-D**) Dissected and stained gonads, showing representative maximal projections. Sperm and oocytes were stained with sp56 and RME-2 antibodies, respectively. Dashed yellow box, region magnified in D’-D’’’’. (**D’-D””**) Sperm and oocyte markers shown individually and merged. Overlap of sperm and oocyte markers in *mpk-1b(Δ)* is highly abnormal. (**E-F**) Box plots of gonad volume and germ cell number for *mpk-1(DT)*, *mpk-1(Δ)*, and *mpk-1b(Δ)*. Gonad volume was calculated using Imaris (see Methods). Germ cells were counted manually using FIJI. Sample sizes are, from left to right, 10, 6, 7. Red asterisk, statistical significance *vs* all other samples (p < 0.01; ANOVA followed by Tukey); ns, not significant.

Intriguingly, the morphologies of *mpk-1(Δ)* and *mpk-1b(Δ)* germlines were dramatically different. To investigate their differences, we stained dissected gonads from wild-type *mpk-1(Δ)* and *mpk-1b(Δ)* adult hermaphrodites with sperm and oocyte markers plus DAPI (Fig. 3B-D). Wildtype germlines were large with proliferating GSCs in the progenitor zone, neatly organized germ cells in the pachytene region, oocyte staining in a single file row of oocytes in the proximal gonad, and sperm staining restricted from oocyte staining (Fig. 3B; Table S1). By contrast, *mpk-1(Δ)* germlines were much smaller, had a disordered pachytene region, and failed to produce gametes (Fig. 3C; Table S1). This *mpk-1(Δ)* germline morphology is similar to that of *mpk-1(ø)* mutants, reported previously^18, 19^. The *mpk-1b(Δ)* germlines had one feature similar to *mpk-1(Δ)* and *mpk-1(ø)*: the *mpk-1b(Δ)* pachytene region was disorganized, indicating a defect in pachytene progression (Figure 3D; Table S1). However, unlike *mpk-1(Δ), mpk-1b(Δ)* germlines were large and appeared similar in size to wild-type (Fig. 3D-E). In addition, most *mpk-1b(Δ)* germlines initiated gamete formation (Fig. 3D). Thus, 95% of *mpk-1b(Δ)* germlines were positive for a sperm marker, and 66% were also positive for an oocyte marker (Table S1). Yet the sperm and oocytes were not arranged normally: cells staining with the sperm marker were not spatially restricted to the proximal end, as in wild-type, but instead they were intermingled with malformed cells staining with the oocyte marker (Fig. 3D’-3D””). We draw two conclusions from this data. First, MPK-1B is required for organization of the pachytene region, but can be dispensable for pachytene exit and the initiation of gametogenesis. Second, MPK-1A can initiate gametogenesis in *mpk-1b(Δ)* mutants, although the gametes are aberrant. Because MPK-1A is not expressed in the germline, its germline effects must be non-autonomous.

We were struck that some *mpk-1b(Δ)* gonad arms appeared larger than wild-type (compare Fig. 3B to D). To probe further, we calculated gonad arm volumes and counted germ nuclei in wild-type, *mpk-1(Δ)*, and *mpk-1b(Δ)* (Fig. 3E-F). As expected, *mpk-1(Δ)* arms were substantially smaller than wild-type by both volume and number of nuclei. The *mpk-1b(Δ)* gonadal arm, on the other hand, had a volume similar to that of the wild-type (Fig. 3E), but number of nuclei were reduced by ~25% compared to wild-type (Fig. 3F). Therefore, *mpk-1b(Δ)* mutants make a germline of comparable size to the wild-type, but with fewer nuclei. Because *mpk-1b(Δ)* retain MPK-1A activity, we infer that somatic MPK-1A promotes germline growth, both in terms of volume and germ cell number.

### Germline MPK-1B does not promote GSC proliferation

We next tested the roles of MPK-1A and MPK-1B in GSC proliferation. To this end, we expressed GFP::MPK-1B in *mpk-1(ø)* (the *ga117* allele^15^) mutants (henceforth *germline::MPK-1B*), using a single-copy transgene driven by the germline-specific *mex-5* promoter^47, 48^. These animals possess MPK-1 activity in the germline due to the MPK-1B transgene, but lack MPK-1 activity in the soma due to the *mpk-1(ø)* mutation; as a result, this strain is effectively a null mutant for MPK-1A. The germline expression of GFP::MPK-1B rescued the *mpk-1(ø)* sterility, but did not rescue *mpk-1(ø)* vulval defects. The animals generated progeny, but without a vulva, could not extrude embryos from the uterus, forcing them to hatch inside their mother (Fig. 4A-D). The vulva defect in *germline::MPK-1B* confirms its lack of somatic MPK-1 activity. The *germline::MPK-1B* and *mpk-1b(Δ)* results are therefore complementary and together, they demonstrate that MPK-1B is sufficient to promote progression through pachytene and form functional gametes. MPK-1B is also necessary for the formation of functional gametes. In addition, the results show that MPK-1B promotes germline function autonomously.

**Figure 4.**
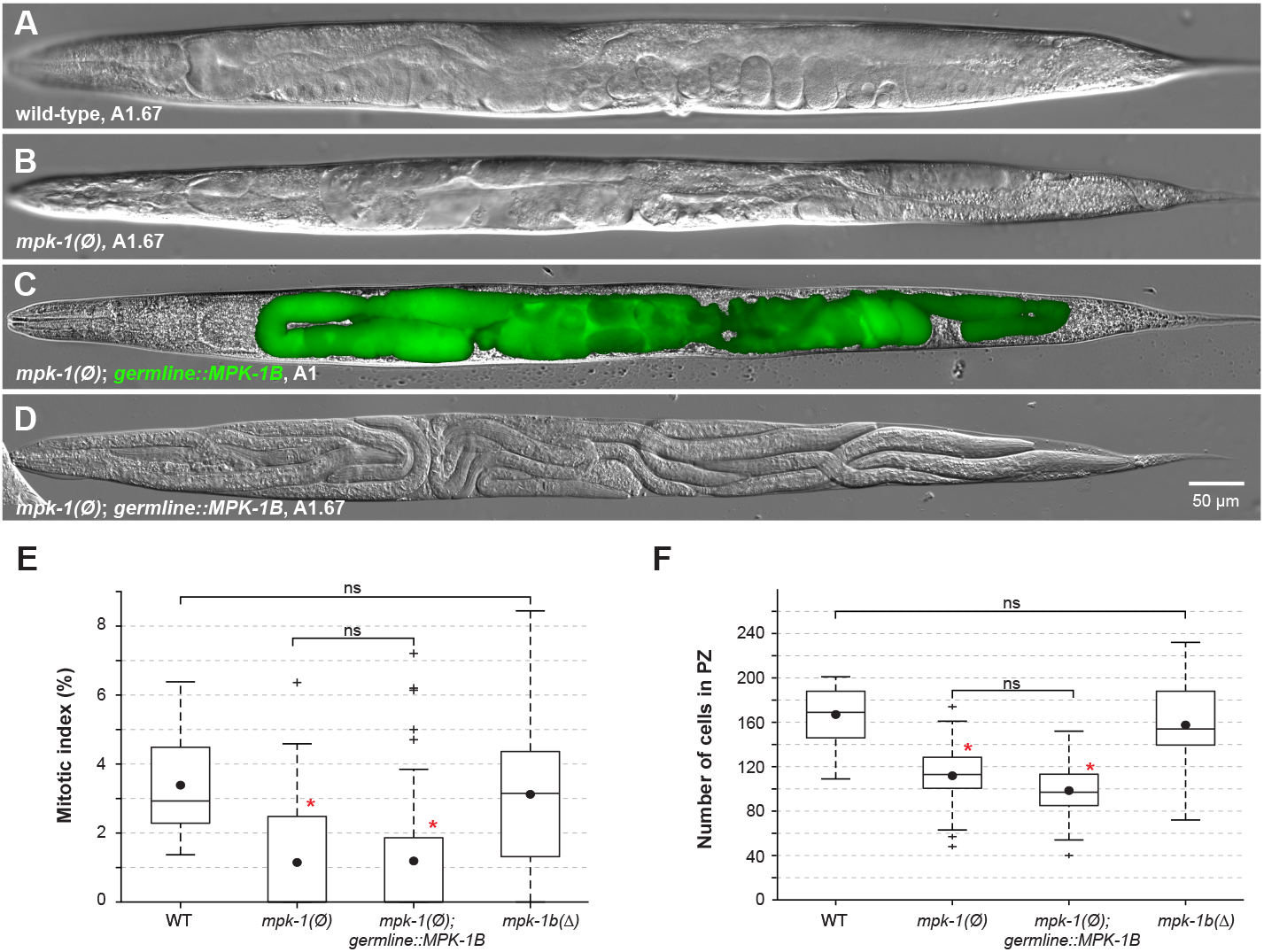
Germline MPK-1B does not promote GSC proliferation. (**A-D**) Representative DIC images of adults of indicated genotypes (full genotypes in Table S4), at either 24 (A1) or 40 (A1.67) hours after the late L4 stage (25°C). GFP::MPK-1B expression is germline-specific and shown as a green overlay in panel C. Anterior is to the left. (**E-F**) Box plots of the progenitor zone mitotic index and germ cell number in A1 hermaphrodites. Dots mark averages. Sample sizes are, from left to right, 14, 37, 39, 32. Red asterisk, statistical significance *vs* wild-type (WT) (p < 0.01; (**E**) Kruskal-Wallis followed by Dunn; (**F**) ANOVA followed by Tukey).

As might be expected for such a small germline, the GSC Mitotic Index in *mpk-1(ø)* mutants is lower than wild-type (Fig. 4E)^18, 21^. The number of germ cells in the Progenitor Zone is also reduced in these mutants (Fig. 4F). Since germline MPK-1B was sufficient to restore fertility in *mpk-1(ø)* mutants, we asked whether it also restored the GSC MI and PZ cell numbers in *mpk-1(ø)* mutants. To this end, we measured MI and PZ cell numbers in *germline::MPK-1B* animals. Strikingly, despite rescuing fertility, germline MPK-1B did not restore either the MI (Fig. 4E) or number of germ cells in the PZ (Fig. 4F). The simple explanation was that MPK-1A must be required in the soma for both GSC MI and PZ size. To test this idea, we scored *mpk-1b(Δ)* mutants for these two traits. As expected, the GSC MI and PZ cell number were indistinguishable in *mpk-1b(Δ)* and wild-type (Fig. 4E-F). We draw several conclusions from these experiments. First, somatic MPK-1A is required for the normal number of adult PZ cells. Second, germline MPK-1B is not required for adults to achieve high GSC proliferation rates; the reduction in total germ cell number found in *mpk-1b(Δ)* germlines is therefore likely due to problems in meiotic progression. Third and most important, the somatic MPK-1A isoform is non-autonomously required to promote the high GSC proliferation typical of young adult hermaphrodites, while germline MPK-1B is dispensable.

### MPK-1A promotes GSC proliferation non-autonomously from the gut and/or somatic gonad

To determine where MPK-1A acts in the soma to promote GSC proliferation, we turned to transgenic arrays driving MPK-1A expression in somatic tissues in the *mpk-1(ø)* background. Because these arrays are efficiently silenced in the germline^49, 50^, expression was limited to somatic tissues. To follow array transmission, we used muscle-expressed mCherry (henceforth *muscle::mCherry*) as a co-transformation marker, which also provided vulva muscle landmarks (Fig. 5A-I, yellow arrows: normal). Our negative control expressed *muscle::mCherry* alone in *mpk-1(ø)* mutants and had no rescuing effect (Fig. 5B). Our positive control carried muscle-expressed mCherry plus GFP::MPK-1A expressed in all somatic cells under control of the *sur-5* promoter (henceforth *soma::MPK-1A*)^51^. The *sur-5* promoter drove strong GFP::MPK-1A expression in the gut, and lower levels in all other somatic cell types (Fig. 5C). As expected, *soma::MPK-1A* rescued vulva formation, albeit not in all animals likely due to mosaicism (Figs. 5A-C, S4; Table S2). Most importantly, *soma::MPK-1A* restored the GSC Mitotic Index to that of *muscle::mCherry* controls (Fig. 5J). The Progenitor Zone cell counts however remained lower than those of controls (Fig. 5K). We do not understand why PZ size was not restored, but transgenic mis-expression may play some role. Consistent with that possibility, *soma::MPK-1A* sometimes induced a multivulva (Table S2), a common sign of MPK-1 hyperactivity^15^. Although able to restore GSC proliferation, *soma::MPK-1A* could not restore fertility (Fig. 5A-C, Table S3), further confirming that germline MPK-1B is essential for germline function (Fig. 3A). Fertility and vulva development were both rescued only when *soma::MPK-1A* and *germline::MPK-1B* were combined into *mpk-1(ø)* mutants (Fig. 5D; Table S3). We conclude that somatic MPK-1A is sufficient to drive the high GSC proliferation typical of young adult hermaphrodites and therefore acts non-autonomously. In addition, the unexpected non-rescue of adult PZ size suggests that this parameter is likely regulated independently from the GSC proliferation rate.

**Figure 5.**
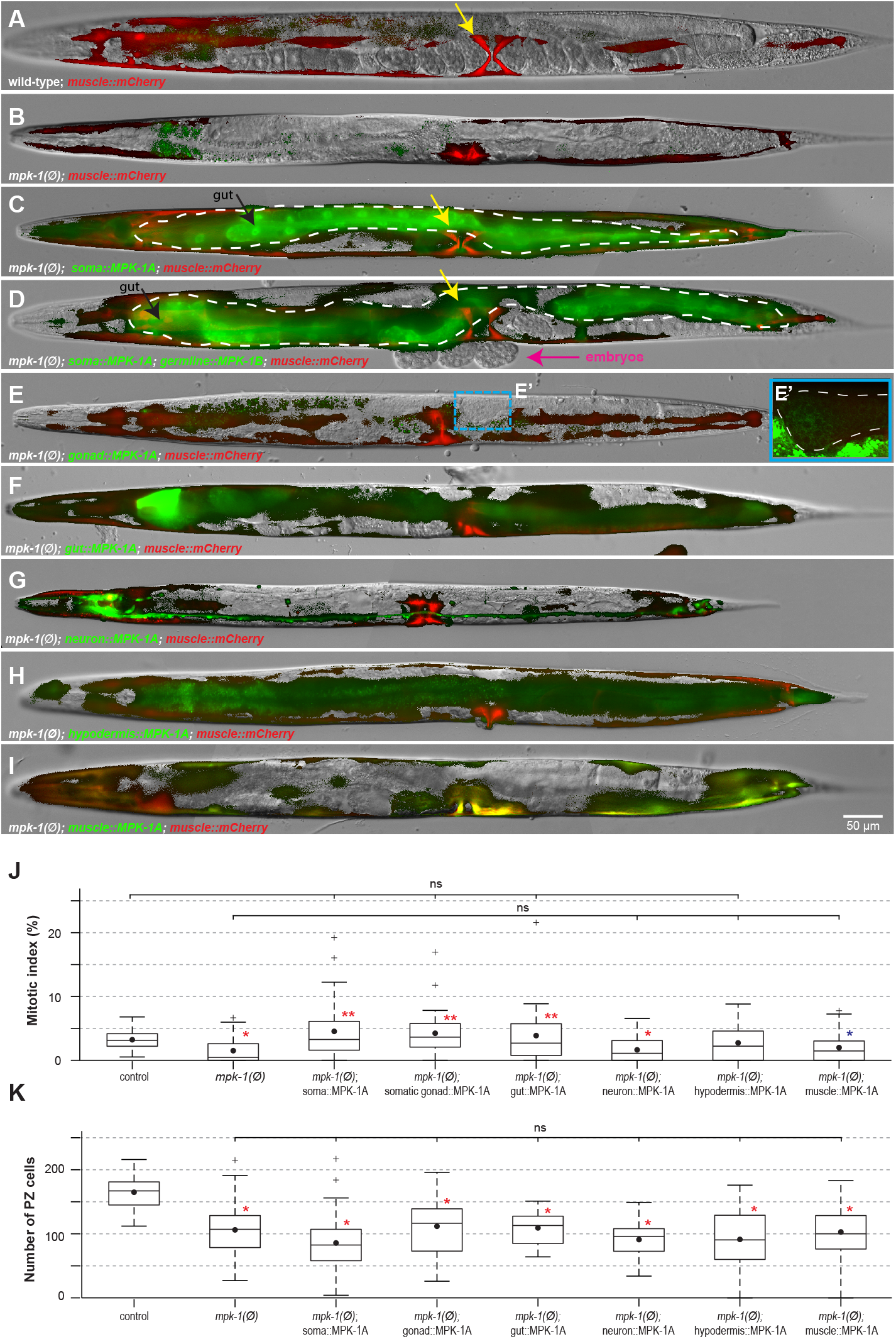
Somatic MPK-1A promotes GSC proliferation non-autonomously. (**A-I**) Representative DIC images of A1 hermaphrodites of indicated genotypes (full genotypes in Table S4), overlaid with fluorescence signals from GFP::MPK-1A and muscle::mCherry. All animals carry *Pmyo-3::mCherry* as an extrachromosomal array marker, and also to assess vulva muscle specification (yellow arrows mark properly specified vulva muscles). Anterior is to the left. Images were not all captured with the same settings, because GFP::MPK-1A levels varied from promoter to promoter, and the aim was to illustrate specificity and pattern of expression. Qualitatively, somatic, neuronal, and muscle promoters drove stronger GFP::MPK-1A expression, while gut, hypodermis, and gonad promoters drove weaker expression. Dashed lines, gut expression in C-D; somatic gonad expression in E’. Notes: For unknown reasons, animals in (D) appeared hypersensitive to tetramisole and laid embryos (pink arrow) upon paralysis. Also in these animals, GFP::MPK-1B (in the germline) was unusually low (compare to Fig. 4C), likely due to co-suppression^73^. Box plots of (**J**) the progenitor zone mitotic index and (**K**) germ cell number in A1 hermaphrodites. Dots mark averages. For each genotype, data from two to three independent lines were pooled together, except for the control, where only one line was analysed. (**J**) Sample sizes are, for each genotype from left to right, 24, 39, 40, 34, 31, 39, 25, 42. Single asterisk, statistical significance *vs* control; two asterisks, *vs mpk-1(ø)* (Red: p < 0.01, blue: p < 0.05; Kruskal-Wallis followed by Dunn). (**K**) Sample sizes are, for each genotype from left to right, 24, 39, 40, 34, 31, 39, 26, 43. Red asterisk, statistical significance *vs* control (p < 0.01; ANOVA, followed by Tukey).

We next used tissue-specific promoters to drive MPK-1A in individual somatic tissues of *mpk-1(ø)* animals. Specifically, we used *rgef-1, dpy-7, elt-7, myo-3*, and *ckb-3* promoters to drive expression in the nervous system, hypodermis (hyp 7), gut, non-pharyngeal muscles and somatic gonad, respectively^52-56^. We achieved high GFP::MPK-1A expression in the nervous system and non-pharyngeal muscles, intermediate levels in the hypodermis and gut, and lower levels in the somatic gonad of young adult hermaphrodites (Fig. 5E-I). Tissue-specific expression of MPK-1A in either the gut or somatic gonad fully rescued the germline Mitotic Index of *mpk-1(ø)* mutants (Fig. 5J). By contrast, expression in the nervous system, non-pharyngeal muscles or hypodermis had no significant effect on the MI (Fig. 5 J). As seen in *soma::MPK-1A* animals, none of the tissue-specific promoters rescued PZ size (Fig. 5K). We conclude that MPK-1A acts either in the gut and/or in the somatic gonad to non-autonomously support the high GSC proliferation that is observed in young adult hermaphrodites.

### MPK-1A promotes GSC proliferation independently of other known roles

In addition to the well-established *mpk-1(ø)* defects in vulval and germline development, we noticed two additional defects. The *mpk-1(ø)* mutants wandered outside the bacterial lawn much more often than wild-type (Fig 6A), and their average body length was 10% longer than wild-type (Fig. 6B). We considered the possibility that one of these defects might be linked to the reduced GSC proliferation of *mpk-1(ø)* mutants. If true, these additional defects should be rescued concurrently with GSC proliferation.

**Figure 6.**
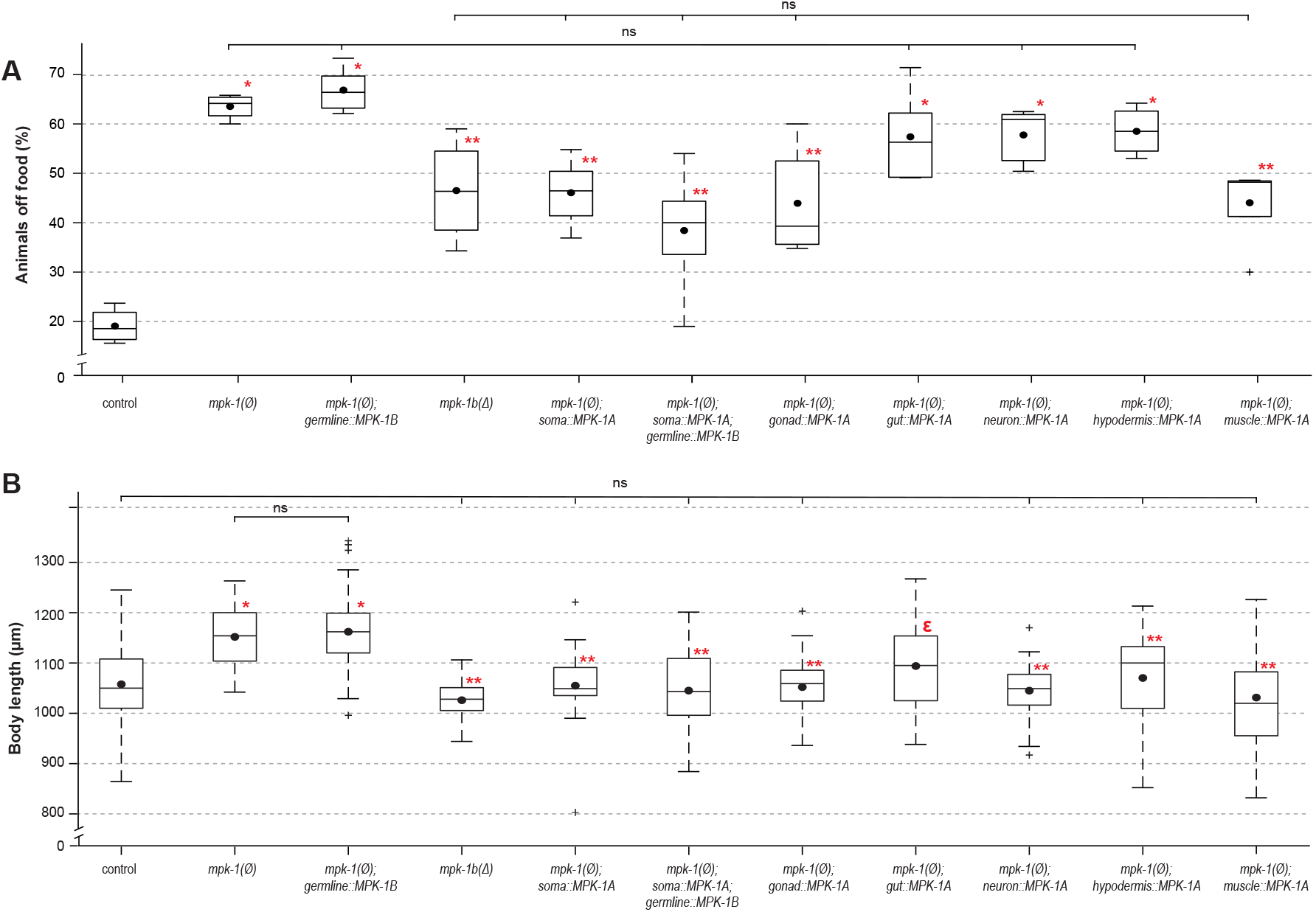
Regulation of food attraction and body length by somatic MPK-1A. Box plots of (**A**) wandering behavior, measured as the percentage of animals found outside of the bacterial lawn and (**B**) body length in adult hermaphrodites of indicated genotypes (full genotypes in Table S4). (**A-B**) Red asterisk, statistical significance *vs* control; two asterisks, *vs* (**A**) both control and *mpk-1(ø)*, or (**B**) *vs mpk-1(ø)* (p < 0.05; One-way ANOVA, followed by Tukey HSD multiple comparisons). ε, Body length of *gut-rescued mpk-1(ø)* was not significantly different from control and from *mpk-1(ø);* it was only significantly different from *mpk-1(ø); muscle::MPK-1A* in pairwise comparisons. Sample sizes are, for each genotype from left to right, (**A**): 4, 4, 6, 4, 6, 7, 5, 6, 5, 4, 5, each with cohorts of 20-30 animals; (**B**): 107, 15, 106, 17, 19, 36, 15, 43, 15, 29, 40.

We therefore asked whether these additional defects were always rescued concurrently with GSC proliferation in the MPK-1 variants we had created in this work. We found that MPK-1A expression in either the somatic gonad or muscles restored the wandering behavior of *mpk-1(ø)* mutants, whereas gut, neuron and hypodermis MPK-1A, or germline MPK-1B, had no noticeable effect (Fig. 6A). For body length, we found that MPK-1A expression in any somatic tissue (except for the gut) was sufficient to prevent excessive elongation, while germline MPK-1B had no significant effect (Fig. 6B). Overall, both defects were restored without a concurrent rescue of GSC proliferation, and conversely, GSC proliferation was restored without a concurrent rescue of the two defects (Figs. 5J-6, 7). We therefore conclude that MPK-1A activity in the animal’s gut and/or somatic gonad promotes GSC proliferation independently from preventing to wander off the food and undergo excessive body elongation.

**Figure 7.**
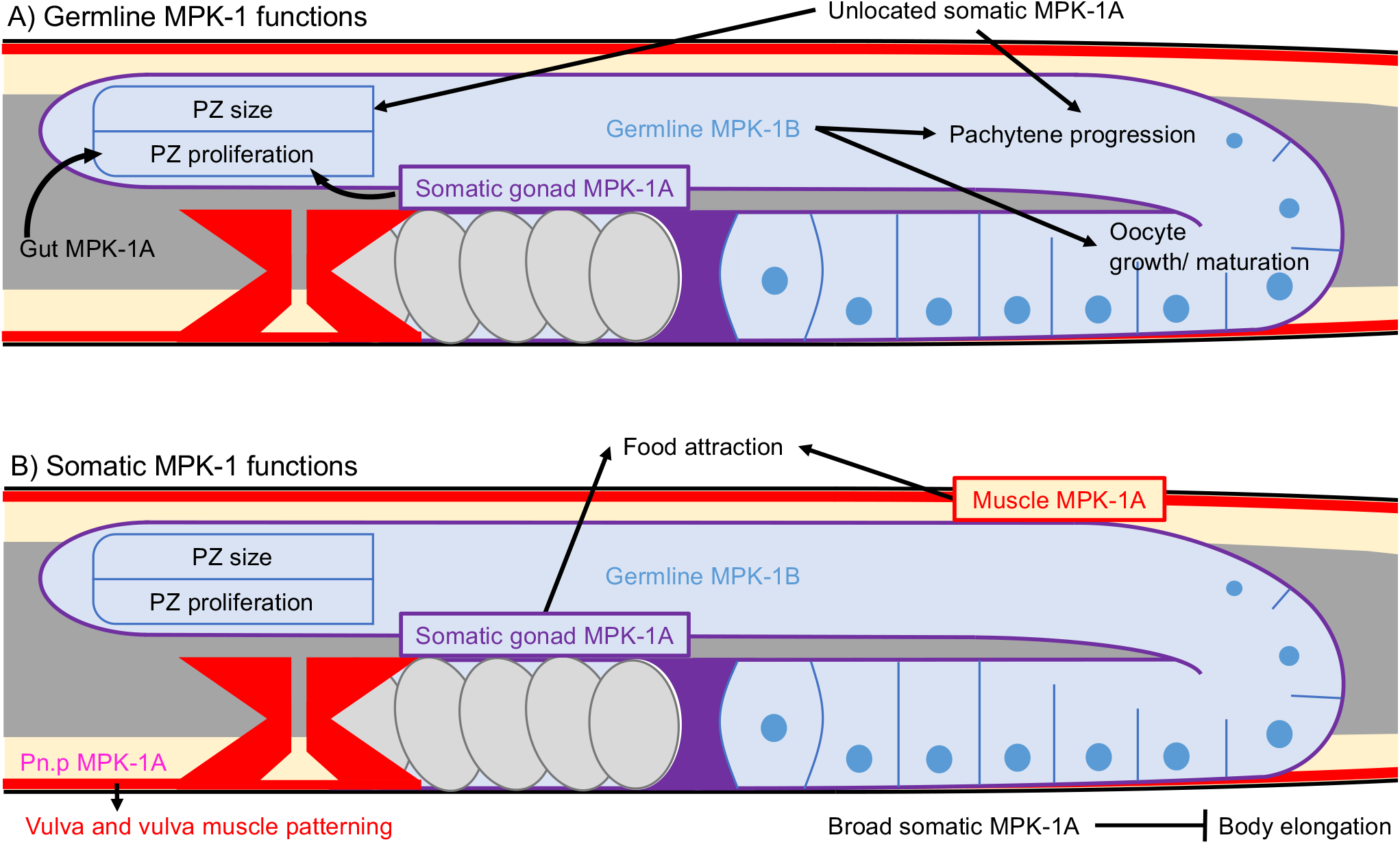
Models for cell autonomous and non-autonomous MPK-1 functions. (**A**) MPK-1 affects the germline both autonomously and non-autonomously. Germline MPK-1B autonomously ensures proper meiotic progression and gametogenesis. Somatic MPK-1A, on the other hand, non-autonomously ensures proper progenitor zone size and proliferation and some, albeit abnormal meiotic progression and gametogenesis. MPK-1A non-autonomously promotes GSC proliferation from the gut and somatic gonad, but its site of action for meiotic progression and gametogenesis remain unknown. (**B**) MPK-1 activity has broad effects on somatic functions. This conclusion is well established for vulval tissues^16^, but this work adds MPK-1 effects on food attraction and body elongation. Somatic MPK-1A is thought to act autonomously in the vulva^16^; it promotes food attraction non-autonomously from the somatic gonad and muscle and regulates body length from any somatic tissue except the gut. (**A-B**) Arrows represent stimulation, and bars, repression. Gut, grey. Somatic gonad, purple. Germline, blue. Muscles, red.

## Discussion

This work advances our understanding of how ERK/MAPK regulates proliferation in the *C. elegans* germline with implications for ERK/MAPK regulation of animal development and human health. In contrast to earlier studies, we detected MPK-1 activity within proliferating germ cells, including germline stem cells (GSCs), but surprisingly, that activity did not correlate with proliferation. Analysis of MPK-1 isoforms led to the answer. The germline isoform, MPK-1B, has no apparent effect on proliferation but instead controls differentiation, whereas the somatic isoform, MPK-1A, acts non-autonomously from the gut or somatic gonad to promote germline proliferation. To our knowledge, this is the first demonstration of a non-autonomous effect of ERK/MAPK on stem cell proliferation within an animal.

Prior to this work, evidence for MPK-1 regulation of germline proliferation was paradoxical. MPK-1 clearly promotes GSC proliferation, as deduced from *mpk-1* null mutants^18, 21^, but active MPK-1 was not found in proliferating germ cells, instead being restricted to meiotic germ cells^18^. To address this paradox, we used a highly sensitive reporter and identified significant MPK-1 activity in proliferating germ cells, though activity is low. However, the level of MPK-1 activity in proliferating germ cells did not correlate with the level of proliferation. Thus, gain-of-function *ras* mutants, *let-60(gf)*, had no detectable MPK-1 activity in the progenitor zone but continued to proliferate normally, while loss of function *fog-1* mutants, had normal levels of MPK-1 activity in the PZ but proliferation was arrested. Therefore, MPK-1 activity in proliferating germ cells does not appear relevant to their proliferation. Future studies will be required to ask if MPK-1 activity plays some other role, perhaps in stem cell genomic integrity, similar to its mammalian counterpart^11^.

The paradox was solved by identifying the functions of ERK/MAPK isoforms, one expressed in the germline and the other in the soma. We created animals that express MPK-1A but not MPK-1B, and vice versa. The MPK-1A-only animals were made with a deletion in the *mpk-1b*-specific exon, and MPK-1B-only animals with a germline-driven transgenic MPK-1B in an *mpk-1* null mutant. Remarkably, germline proliferation was essentially normal in MPK-1A-only animals but not in MPK-1B-only animals. Therefore, somatic MPK-1A is the key driver of germline proliferation and must act non-autonomously. However, either somatic MPK-1A or germline MPK-1B can promote more extensive differentiation than found in *mpk-1(ø)* mutants. Germline MPK-1B is sufficient to ensure formation of fully functional gametes at least in some animals, while somatic MPK-1A provides a non-autonomous boost on differentiation (Fig. 7A). In the absence of MPK-1B, that MPK-1A boost is sufficient for pachytene exit and initiation of gametogenesis but gametes are not functional.

The *C. elegans* ERK/MAPK non-autonomous effect on germline proliferation relies on expression in the gut and/or somatic gonad. Intriguingly, gap junctions connect gut and somatic gonad as well as somatic gonad and germ cells, and moreover, those gap junctions are essential for germline proliferation^57^. Though speculative, an attractive idea is that somatic MPK-1A uses those gap junctions in some manner to stimulate germline proliferation. In mice, ERK/MAPK phosphorylation of connexins modulates the opening of gap junction during epidermal wound healing^58, 59^. By analogy, ERK/MPK-1 could modulate gut and/or gonadal gap junctions to allow unidentified proliferation-stimulatory small molecules to reach the germline. The identity of such molecules remains unknown. However, one possibility is uridine or thymidine. That possibility emerges from a study of cytidine deaminases (ccd), enzymes that make uridine and thymidine from cytidine and that are required for normal germline proliferation^59^. Remarkably, the defective germline proliferation in *ccd* mutants is rescued by expression of cytidine deaminase in the gut or somatic gonad^60^, solidifying the gut-gonad-germline axis of proliferation control. Yet a direct relationship between MPK-1A and germline uridine/thymidine levels remains highly speculative.

Discovery of the non-autonomous role for ERK/MAPK in germline proliferation has implications for the homeostatic regulation of germline proliferation in response to oocyte accumulation. That regulation is reliant on MPK-1 inhibition locally within the affected gonadal arm, which includes both somatic and germline cells. We suggest that homeostatic inhibition of MPK-1 signaling might occur in the gut or somatic gonad given that MPK-1 functions there to promote proliferation (this work). We favor the somatic gonad as the likely site for MPK-1 inhibition during homeostasis. Each gonadal arm is embraced independently by somatic gonadal sheath cells, which connect to the germline via gap junctions. By contrast, the gut runs the length of the body cavity, neighbors both gonadal arms and is likely connected to each equally via gap junctions. Because homeostatic control of proliferation occurs broadly in the animal kingdom and because regulators of stem cell proliferation and homeostasis are highly conserved (ERK/MAPK and IIS, see Introduction), these regulators may similarly combine to orchestrate stem cell proliferation rates in diverse stem cell populations. Consistent with this, GSC proliferation in *Drosophila* is stimulated by nutrient uptake via IIS signaling^33^, while homeostatic regulation of intestinal stem cells occurs through ERK/MAPK regulation^61^.

The non-autonomous role for ERK/MAPK in the regulation of germline proliferation has important implications for understanding and perhaps treating cancer. ERK/MAPK is upregulated in many human cancers^1-4^, though its primary role in embryonic stem cells centers on differentiation^7-10^. Importantly, tumours are heterogeneous and only cancer stem cells generate a new tumour upon transplantation^62, 63^. The bulk of the tumour therefore likely consists of the progeny of cancer stem cells, which are in various states of differentiation. Cancer stem cells are thought to develop from non-cancerous stem cells as a result of replication errors^64-66^ and thus retain stem cell character. As such, cancer stem cells may depend on ERK/MAPK activity in neighboring tissues or cells to ensure their high proliferation – similar to *C. elegans* somatic ERK/MAPK ensuring high GSC proliferation. If this is the case, chemotherapy that lowers ERK/MAPK activity might either promote quiescence in cancer stem cells non-autonomously, and/or suppress tumour growth by autonomously inhibiting the cancer stem cell’s differentiated progeny. The specificity and effectiveness of chemotherapy may therefore be increased by targeting ERK/MAPK in the tissue supporting cancer stem cell proliferation in addition to inhibiting ERK/MAPK in the cancer stem cell’s differentiated progeny.

## Methods

### *C. elegans* genetics

Animals were maintained on standard NGM plates seeded with *E. coli* (OP50) and the Bristol isolate (N2) was used as wild-type throughout^67^. All animals were scored as young adults with the following specifics. For Figs. 1, 4-5 and S1-S2, animals were maintained at 15°C, synchronized by picking late L4 stage larvae to a new plate^68^, and upshifted to 25°C for 24 hours (unless otherwise specified) before they were harvested for assaying. For Figs. 2-3 and S3, animals were maintained at 20°C and were analyzed at L4 + 24 hours. For Figs 6 and S4, animals were raised at 25°C from the L1 stage. All strains, alleles, transgenes and rearrangements used are listed in Table S4.

### Plasmid constructions, transgenics and genome editing

We used the Gibson method^69^ for all plasmid assembly, except for pUMP5, which was generated by T/A cloning of an RT-PCR product into pMR377, a modified pKSII-based vector, after opening it with *XcmI* to generate T-overhangs (a kind gift from Shaolin Li). The source DNA and primers that were used to generate all plasmids, as well as their microinjection concentrations, are found in Table S5. Extra-chromosomal arrays were generated by regular microinjections at a total concentration of 150-200 ng/μL, using pKSII as a filler DNA and pCFJ104[*Pmyo-3::mCherry*] (5 ng/μL) as a co-injection marker^48, 49^. For the lines harboring pPOM5-9 altogether, each plasmid was injected at 15 ng/μL, in a single mix.

For single-copy insertion (*i.e. narSi2*), pNAR3 was co-injected with pDD122 in *unc-119(ed3)* animals for CRISPR/Cas9-mediated integration at ttTi5605 on LG II (+0.77)^47^. A single line was obtained from > 200 microinjections.

For editing the *mpk-1* locus, CRISPR/Cas9 was used according to previously described methods^70, 71^. Repair templates and crRNAs are listed in Table S6. To create the *mpk-1(DT)*, an intermediate V5-tagged strain was first generated. The 2xOLLAS tag was subsequently inserted into the V5-tagged strain to create *mpk-1(DT)*. The *mpk-1(DT)* strain was used as the starting point to generate *mpk-1b(Δ)*, but the V5-tagged strain was used to generate *mpk-1(Δ)*. All CRISPR created *mpk-1* strains were outcrossed to N2 two times. The *mpk-1b(Δ)* and *mpk-1(Δ)* alleles were maintained over the *qC1[qIs26]* balancer. Full genotypes are listed in Table S4.

### Quantification of germline MPK-1 activity

Following bleach synchronisation, all worms were grown at 25°C until L4 + 24 hours (A1). Animals were collected and paralyzed in 4.15 mM (0.1 %) Tetramisole in M9 buffer on a coverslip that was flipped onto a 3 % agarose pad and sealed using VALAP (1:1:1 Vaseline, lanolin and paraffin). A Leica confocal microscope TCS SP8 (Leica Microsystems) with a HC PL APO CS2 40x/1.30 numerical aperture oil objective was used for image collection. Bidirectional scanning at 400 Hz, in sequential mode, combined with a 0.8 μm Z-stack step was used to image each gonad. Only the anterior gonad arm of each worm was acquired. ERK-nKTR::GFP was acquired using a 488 nm solid-state laser at 10 % intensity, for all strains, except *fog-1(ø)*, which was at 15 % intensity. PMT gate was set at 673 gain and at 559.5±21.5 nm for all strains. *H2B::mCherry* was acquired using a 552 nm solid-state laser at 5 % intensity using a HyD gate at 50 % gain and at 610±21 nm.

Image processing and data analysis were done with Fiji. For the PZ, cells were grouped based on distance from the distal tip (1-5, 6-10, 11-15, and 1-15). Five cells were analysed per gonad, distributed one per three cell diameter regions. For the pachytene region, five cells per gonad were randomly chosen. For oocytes, five cells per gonad were analysed starting from the - 2 oocyte to avoid sperm-activated oocytes. Variability in the *H2B::mCherry* intensities between germ cells and across germline regions (see Fig. S1) prevented us from using the previously published quantification method, developed for vulva precursor cells^43^. Thus, for all germ cells, the mean GFP fluorescence signal intensity was measured for three randomly-chosen circular cytoplasmic areas (0.5 μm for GSCs and pachytene; 4 μm for oocytes) and for the whole nucleus. Cell selection and cytoplasmic area selections were made using the *mCherry* channel to avoid introducing any user-bias after seeing the GFP channel. The three cytoplasmic GFP mean intensity measures were averaged and divided by the single mean nuclear intensity measurement to obtain an MPK-1 activity index for each cell. Five cells of each types were averaged for all strains. For each genotype, the ERK-nKTR activity index for each region was first normalized to its ERK-nKTR(AAA) control, then to the wild-type ERK-nKTR(AAA) control for comparison across different genotypes. As germline autofluorescence accounted for less than 1% of the GFP signal for all samples, background subtractions were omitted.

### GSC pool size and mitotic index

GSC pool size and MIs of young adult hermaphrodites were evaluated as previously described^21, 27^. For every genotype for which we had previously published an A1 MI result (N2, *fog-1(ø)*, and *mpk-1(ø)*)^21, 27^, we did not detect a significant difference between the newer and older datasets (P > 0.05; Kruskall-Wallis rank sum test).

### Immunostaining and fluorescence quantification

#### Sperm/oocyte staining

Briefly, worms raised at 20°C were picked as mid-L4s to a fresh plate, 24 hours prior to staining. Hermaphrodites were anesthetised in 0.25 μM levamisole in PBST (1xPBS + 0.1% Tween20). Gonads were collected and fixed in 3.7% formaldehyde in PBST for 15 minutes while rocking at room temperature (RT). After washing in 1 mL PBST, gonads were permeabilized in PBST+0.1% Triton-X and incubated for 10 minutes, rocking at RT. Gonads were washed 3x 10 minutes in PBST and blocked in PBST+0.5%BSA (block) for 1 hour. After the block was removed, samples were incubated overnight at 4 °C with primary antibodies diluted in the block solution, 1:200 sperm marker mouse α-sp56 and 1: 500 oocyte marker rabbit α-RME-2. Primary antibodies were removed and gonads were washed 3x 10 minutes in PBST. 100 μL of block containing DAPI (1 μg/mL), α-mouse alexa647 and α-rabbit alexa555 secondary antibody (1:1000 each) was added and gonads incubated in the dark, rotating for 2 hours at RT. Gonads were washed 3x 10 minutes in PBST in the dark at RT. Gonads were mounted in 18 μL Prolong Glass antifade (ThermoFisher, P36984) and sealed with VALAP. Samples were kept at −20°C until imaged.

#### MPK-1 staining

Hermaphrodites were staged, anesthetized and dissected as described for sperm/oocyte staining. Gonads were fixed in 2% paraformaldehyde (PFA) in 100 nM pH 7.2 K_2_PO_4_ for 30 minutes, rocking at RT. Gonads were washed 2x in PBST: first wash quick and second wash for 5 minutes at RT. Gonads were fixed in methanol for 30 minutes at −20 °C. Gonads were washed 2x, following the same procedure as after the PFA fix. Gonads were blocked for 1 hour, rocking at RT. Primary antibodies—mouse α-V5 (Bio-Rad), rat α-OLLAS (Novis, NBP1-06713), rabbit α-ERK (Santa Cruz Biotechnology, Sc94)—were diluted 1:1000, 1:200, 1:1000 in blocking solution, respectively. Gonads incubated with primary antibodies overnight at 4 °C. Gonads were then washed 2x PBST quickly and 2x PBST for 10 minutes. Secondary α-mouse alexa555, α-rat alexa647, and α-rabbit alexa488 antibodies, and DAPI (1 μg/mL) were all diluted 1:1000 in block solution and incubated for 2 hours, rocking in the dark at RT. Secondary antibodies were removed and gonads were washed 4x in PBST—2x quickly and 2x for 10 minutes in the dark. Gonads were mounted in 18 μL Prolong Glass antifade, sealed with VALAP, and stored at −20°C until imaged.

#### *DAPI* staining

Staged L4+24 hr hermaphrodites were dissected in PBST+0.25 μM levamisole. Gonads were collected and fixed in 2% pFA for 10 minutes, rocking at RT. The fixation solution was removed; samples were washed once with PBST. The gonads were permeabilized in PBST+0.5%BSA+0.1%Triton-X for 10 minutes, rocking at RT. The permeabilization solution was removed and gonads were incubated with DAPI (1 μg/mL) in PBST for 30 minutes, rocking in the dark at RT. Gonads were washed 3x in PBST for 10 minutes in the dark at RT. After the washes, gonads were mounted in 10 μL Vectashield (Vector laboratories) and sealed with VALAP. Samples were kept at 4°C until imaged.

### Image acquisition

For Figs. 2-3, Fig. S3, all images were taken using a Leica SP8 confocal microscope using a 40x oil objective (NA 1.3) with 1.5 zoom and 0.30 μm z-step. For fluorescence quantification (Fig. S3E-G), images of the distal gonad were taken first (progenitor zone through mid-pachytene region). Fluorophores alexa 488, alexa 555, alexa 647, and DAPI were excited at 488 nm, 561 nm, 633 nm, and 105 nm respectively; emissions were collected at 510-540 nm, 562-600 nm, 650-700 nm, and 425-490 nm, respectively.

The mosaic merge function of the Leica Lightening software package was used to generate Figs. 2-3 images. For all tile scanned germlines, a custom region of interest was drawn around the tissue; images were taken using a 1024×1024 window. All tiles covering the entire germline tissue were merged into one image during acquisition using default settings. We did not quantify fluorescence of the mosaic merged images because the fluorescence intensity was smoothed at tile junction points.

For Figs. 3-4, differential interference contrast (DIC) and epifluorescence images were acquired every micron using a Plan-Apochromat 20x dry objective (NA 0.8) mounted on an inverted Zeiss Axio Observer.Z1. Images were stitched and deconvolved using the Zen software, and straightened using ImageJ. Epifluorescence signals were overlaid to the DIC images using Photoshop CS6.

### Gonad volume quantification and germ cell counts

DAPI stained mosaic-merged gonads were imported into Imaris (version 9.3.1) using the software’s file converter version 9.5. Using the surfaces function, outlines of the gonads were manually drawn around every other z plane throughout the stack. Afterwards, the “create surface” tool made a volume representation based on the manually drawn outlines. Then the volume was calculated in the detailed statistics tab in the surfaces menu.

The same images were used to calculate both gonad volume germ cell numbers. Using the multipoint tool in FIJI, germ cells were manually counted from the distal end to the loop.

### MPK-1 protein quantification

Fluorescence intensity was measured in FIJI using previously described methods^25, 72^. OLLAS and V5 intensities were normalized to the N2 background. ERK intensity was normalized to the *mpk-1b(Δ)* background because *mpk-1b* was previously shown to be the main germline isoform^19^. Intensity plots were generated by importing FIJI data into MATLAB using the shadedErrorBar function (Rob Campbell, www.GitHub.com, 2020).

### *E. coli* attraction

Animals were raised at 25°C until the late L4 stage and picked, in cohorts of 20-30 individuals, to the center of a 6mm NGM dish, pre-seeded with a 40 uL drop of *E. coli* (OP50) overnight culture. Plates were imaged every half-hour during hours 22-24 post-L4 (until A1). The percentage of worms on food were counted at each time point and averaged over all time points for each plate. This assay was repeated at least 4 times for each genotype.

### Body length measurements

Animals were raised at 25°C until they reached A1, paralysed with tetramizole in M9 buffer, and mounted on a 3% agarose pad. Whole animals were acquired using a 10X objective and measured using ImageJ.

### Statistics

For parametric datasets, the one-way ANOVA was used, and followed by Tukey multiple comparisons. For non-parametric datasets, the Kruskal-Wallis test was used, and followed by Dunn multiple comparisons, adjusted according to the family-wide error rate procedure of Holm, and then by the false discovery rate procedure of Benjamini-Hochberg.

## Acknowledgements

We thank Shaolin Li for sharing reagents and precious advice, Claire de la Cova, Iva Greenwald and Alex Hajnal for sharing strains, Jean-Claude Labbé and Geneviève Pépin for critically reading the manuscript, and Wormbase for its essential role in *C. elegans* research. This work was funded by grants from the UQTR Foundation, the Fonds de Recherche du Québec – Nature et Technologies (2018-NC-205752), the Fonds de Recherche du Québec – Santé (#265445), the Canada Natural Science and Engineering Research Council (RGPIN-2019-06863, RGPAS-2019-00017, DGECR-2019-00326), and the Canada Foundation for Innovation (#36916) to PN. PN is a Junior 1 Bursary Scholar (#252405) of the Fonds de Recherche du Québec – Santé and holds a UQTR research chair. SR-T was supported by the National Science Foundation Graduate Research Fellowship under grant No. (DGE-1256259) and the NIH Predoctoral training grant in Genetics 5T32GM007133. JK was an HHMI Investigator and is now supported by NIH R01 GM134119. Any opinion, findings, and conclusions or recommendations expressed in this material are those of the authors(s) and do not necessarily reflect the views of the National Science Foundation. Some strains were provided by the CGC, which is funded by NIH Office of Research Infrastructure Programs (P40 OD010440).

## Author contribution

BD: Figures 1A-D, S1-2, with guidance by PN.

SR-T: Figures 2-3, S3 and Tables S1, S6, including the generation of all related strains, with guidance by JK and PN. Participated in study design, manuscript drafting and editing.

POM: pPOM5-9 construction, Figures 6, S4 and Table S2 with guidance by PN.

XL: pXA2 construction, with guidance by PN.

AADB: Figure 1E (*fog-1, let-60gf*), with guidance by PN.

VR: Figure 1E (*lin-3*) and Figure 7 drawing, with guidance by PN.

YC: Helped POM with some aspects of Figures 6 and Table S2, including with data analysis.

JK: Manuscript drafting and editing.

PN: Figures 3, 4 (with help from POM for 4A-I), Tables S2-5, in addition to generating some strains for Figure 1, general study design, plasmid design, manuscript drafting and editing.

All authors commented on the manuscript and approved its final version.

## Materials & Correspondence

Correspondence and material requests should be addressed to

## Supplementary material

### Legends to Supplementary Figures

**Figure S1. Germline ERK-nKTR analysis.** (**A-F**) Representative confocal images and (**A’-F’**) closeups of the (**A, C, E**) ERK-nKTR(AAA)::GFP and (**B, D, F**) ERK-nKTR::GFP fluorescence signals in the (**A-B**) Progenitor Zone, (**C-D**) pachytene, and (**E-F**) oocytes in an A1 hermaphrodite of a wild-type background. A different focal plane of the same animal is shown in (**A, C, E**) and (**B, D, F**). Red, *H2B::mCherry;* green, (**A, C, E**) *ERK-nKTR(AAA);* (**B, D, F**) *ERK-nKTR*. A large dotted circle delineates one nucleus that is in focus for each zone. For each delineated nucleus, 3 smaller dotted circles were randomly placed (using the red channel only) around the nucleus for cytoplasmic green channel intensity sampling. (**G, I-K**) Raw cytoplasmic:nuclear GFP fluorescence intensity ratios for both *ERK-nKTR(AAA)* and *ERK-nKTR* in the PZ, pachytene and oocytes in (**G**) wild-type, (**I**) *lin-3(ø)*, (**J**)*fog-1(ø)*, and (**K**) *let-60(ga89)gf* backgrounds. (**H**) Data from (**G**) transformed to represent MPK-1 activity as a percentage of MPK-1 activity in oocytes. (**L**) MPK-1 is similarly active in the pachytene region of control and *let-60(gf)* animals, but not in sterile *fog-1(ø)* and *lin-3(ø)* mutants. (**M**) MPK-1 is similarly active in the oocytes of control and *let-60(gf)* animals, but is significantly lower in unfertilized oocytes of *fog-1(ø)* mutants. (**G-M**) Error bars, standard deviation. Sample sizes, as in Fig. 1 E-F. Black asterisks represent statistical significance in pairwise comparisons; red asterisk, *vs* the *nKTR(AAA)* baseline control (P < 0.01; ANOVA followed by Tukey); ns, not significant.

**Figure S2. Germline phenotypes of selected mutants.** Representative confocal sections of A1 hermaphrodite gonads visualized with *arSi12[ERK-nKTR::GFP; mCherry::H2B]*. Non-germline tissues were masked to emphasize the germline. Dashed lines delineate the oocyte region. (**A**) Wild-type animal. Germline is normal. (**B**) *lin-3(ø)* mutant. Oocytes are endomitotic. (**C**) *fog-1(ø)* mutant. Arrested oocytes are stacked. (**D**) *let-60(gf)* mutant. Oocytes are smaller than normal and disorganized.

**Figure S3. Somatic and germline specificity of MPK-1 isoforms.** (**A-D**) Representative maximal projection MPK-1 staining of distal germlines and/or somatic tissues of wild-type, *mpk-s1(DT)* and *mpk-1b(Δ)* A1 hermaphrodites. Germline, white dashed boundary; somatic, orange dashed boundary. Left, anti-ERK antibodies detect MPK-1A/B in both soma and germline, but not in the *mpk-1b(Δ)* germline. Middle left, anti-OLLAS stains MPK-1(DT) in both germline and gut, no signal is detected in wild-type and *mpk-1b(Δ)* germlines; middle right, anti-V5 detects MPK-1B in the *mpk-1(DT)* germline, but not in somatic tissue (also see Fig. S2), no signal is detected in wild-type and *mpk-1b(Δ)* germlines; right, DAPI. (**E-G**) MPK-1 fluorescence intensity quantification of α-ERK (MPK-1A/B, green), α-OLLAS (MPK-1A/B, yellow), and α-V5 (MPK-1B, red) as a function of position within the distal germline (distal end through to mid pachytene region). See methods for quantification details. Thick colored line is the mean intensity value at each point along the germline. Standard error is represented by shading around the mean.

**Figure S4. Vulva defects in *mpk-1* larvae and rescue by MPK-1A.** (**A-H**) Representative DIC images from L4 larvae of the indicated genotypes. Only *soma::MPK-1A* fully rescued vulva development. (**G**) We noticed that a small number of *Pdpy-7::GFP::MPK-1A* animals appeared to form a partial invagination (as in this example), resulting in a protruding vulva (Pvul) phenotype in the adult (see Fig. 4H). This is likely due to leaky expression from the *Pdpy-7* promoter (reputed as Hyp 7-specific) into the Pn.p cells, which are also part of the hypodermal lineage.

### Supplementary Tables

**Table S1.** Frequency of germline defects in *mpk-1* mutants.

| Genotype | n | Sperm marker<br>(% positive) | Oocyte marker<br>(% positive) | Pachytene region<br>(% normal) |
| --- | --- | --- | --- | --- |
| Wild-type | 23 | 100 | 100 | 100 |
| *mpk-1b*($\Delta$) | 23 | 78 | 43 | 0 |
| *mpk-1*($\Delta$) | 22 | 0 | 0 | 0 |

**Table S2.** Vulva muscle features in *mpk-1(ø)* mutants carrying tissue-specific transgenes.

| Genotype* | Sample size | Transgene lost from vulva muscles (%) | Normal vulva muscles<sup>J</sup> (%) | Abnormal vulva muscle shape (%) | Muv with abnormal vulva muscles (%) |
| --- | --- | --- | --- | --- | --- |
| Control | 57 | 0 | 100 | 0 | 0 |
| *mpk-1*(∅) | 64 | 9.4 | 0 | 90.6 | 0 |
| *mpk-1*(∅); *soma::MPK-1A* | 130 | 6.2 | 49.2 | 40.8 | 3.8 |
| *mpk-1*(∅); *gonad::MPK-1A* | 84 | 11.9 | 0 | 86.9 | 1.2 |
| *mpk-1*(∅); *gut::MPK-1A* | 94 | 8.5 | 0 | 91.5 | 0 |
| *mpk-1*(∅); *muscle::MPK-1A* | 63 | 14.3 | 0 | 85.7 | 0 |
| *mpk-1*(∅); *hypodermis::MPK-1A* | 47 | 6.4 | 0 | 93.6 | 0 |
| *mpk-1*(∅); *neuron::MPK-1A* | 151 | 5.3 | 0 | 94.7 | 0 |
| *mpk-1*(∅); *5 tissues::MPK-1A*<sup>◇</sup> | 36 | 16.7 | 0 | 83.3 | 0 |

\*, Except for the control, all animals are non-Rol mCherry+ progeny from parent of genotype *mpk1(gal117)/qC1[rol-6(gf); lag-2p::GFP]*. All were scored at A1 stage. All animals contain the *Pmyo-3::mCherry* extrachromosomal array, a marker of non-pharyngeal muscles, including vulva muscles. None of the transgenes rescued fertility.
<sup>J</sup>, Vulva muscle formation is rescued by ubiquitous somatic MPK-1A expression, but not by expression in the somatic gonad, gut, hypodermis, neurons and/or muscles. These results are consistent with MPK-1A acting in epithelial Pn.p cells to specify vulva divisions autonomously<sup>16</sup>, and shape vulva muscles non-autonomously.
<sup>◇</sup>, Plasmids for gonad, gut, muscle, hypodermis and neuron MPK-1A expression (pPOM5-9) were injected together at 20 ng/ul each.

**Table S3.** Rescue of *mpk-1(ø)* fertility.

| Genotype | Sample size | Average brood size<br>± standard deviation | Maximal brood size |
| --- | --- | --- | --- |
| Wild-type | 10 | 281 ± 42 | 336 |
| *mpk-1*(∅) | > 1000 | 0 | 0 |
| *soma::MPK-1A* | 130 | 0 | 0 |
| *germline::MPK-1B* | 20 | 12 ± 6 | 24 |
| *soma::MPK-1A; germline::MPK-1B* | 24 | 56 ± 24 | 108 |

**Table S4.**
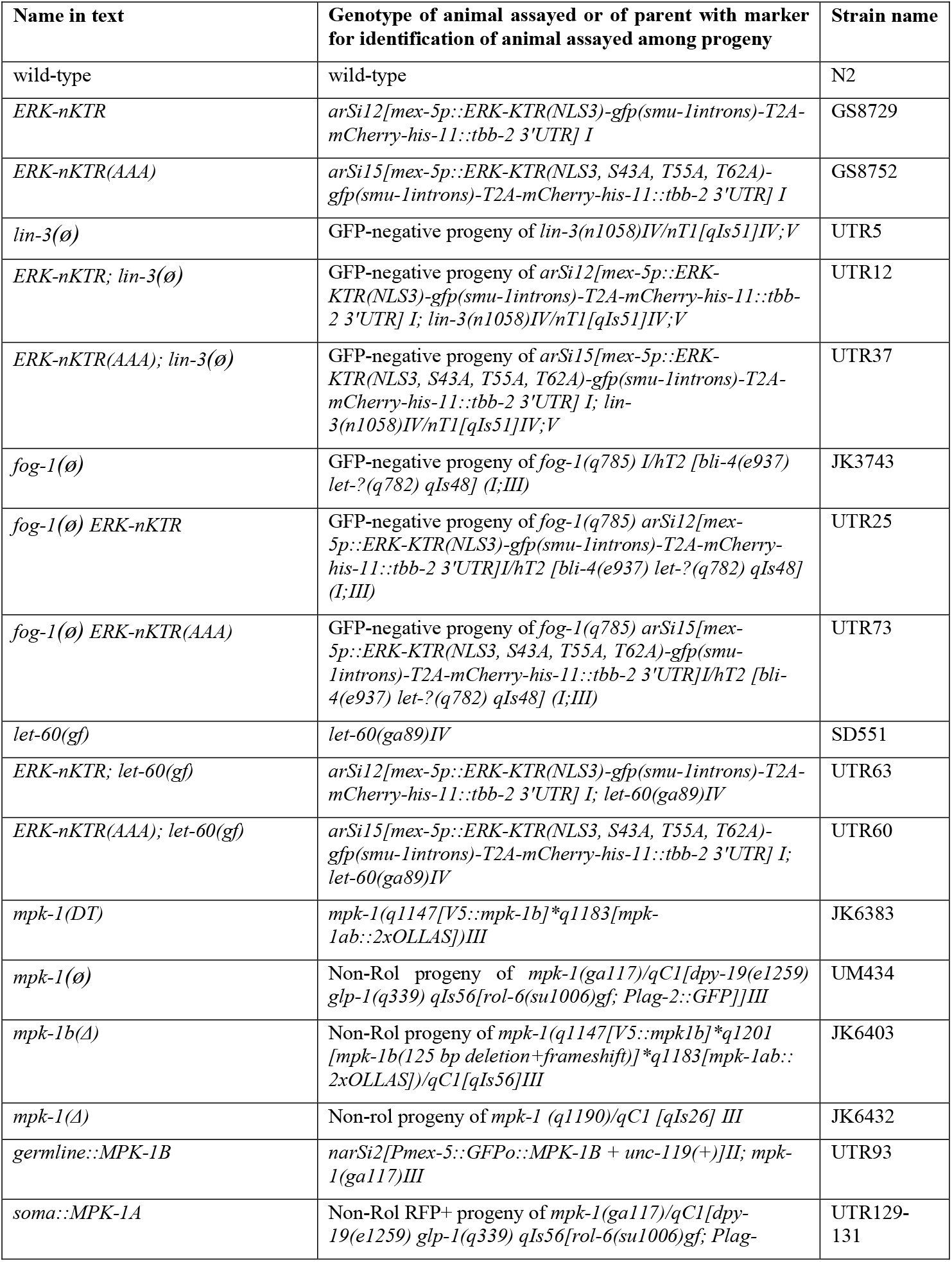
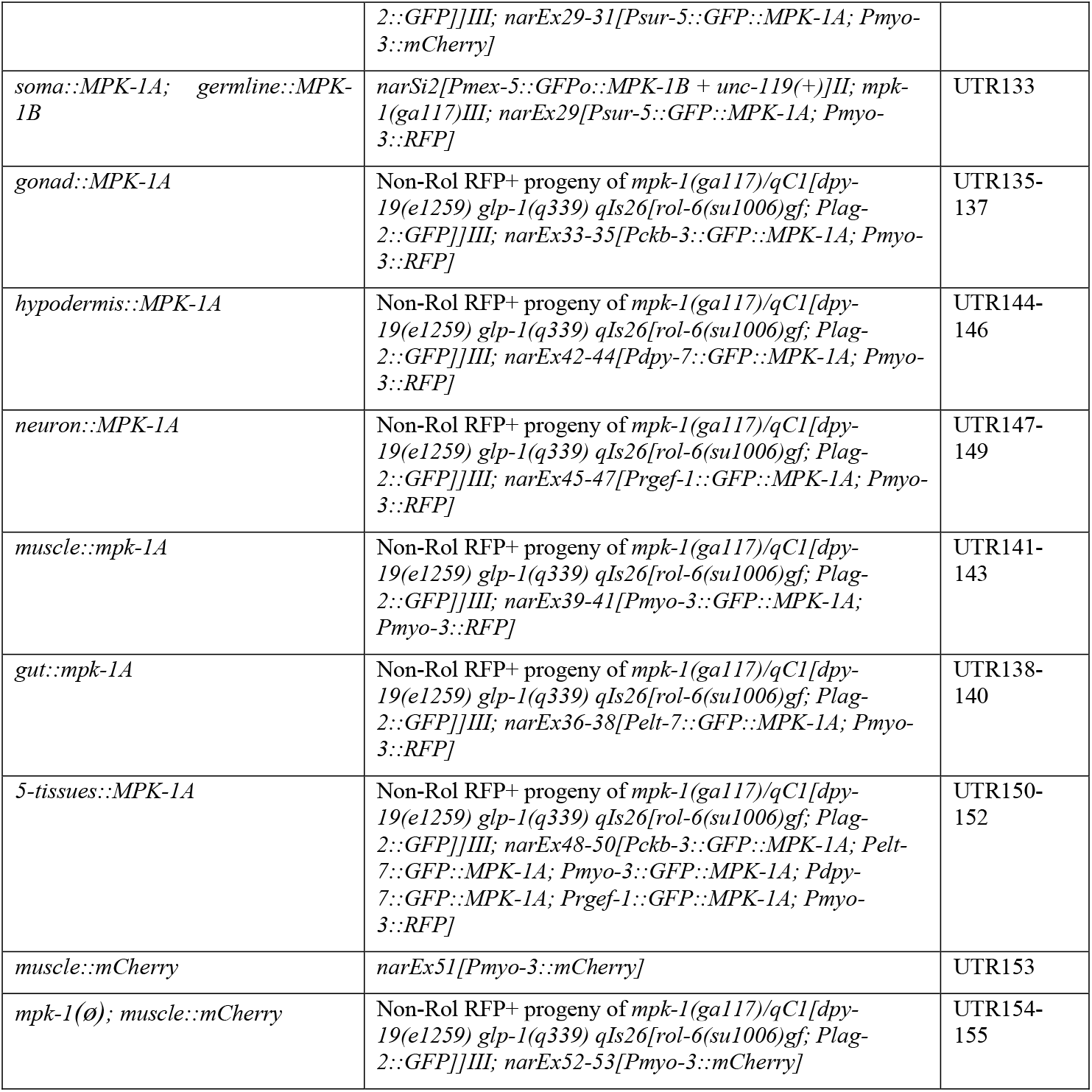
Strains, alleles, transgenes and rearrangements used in this work.

**Table S5.**
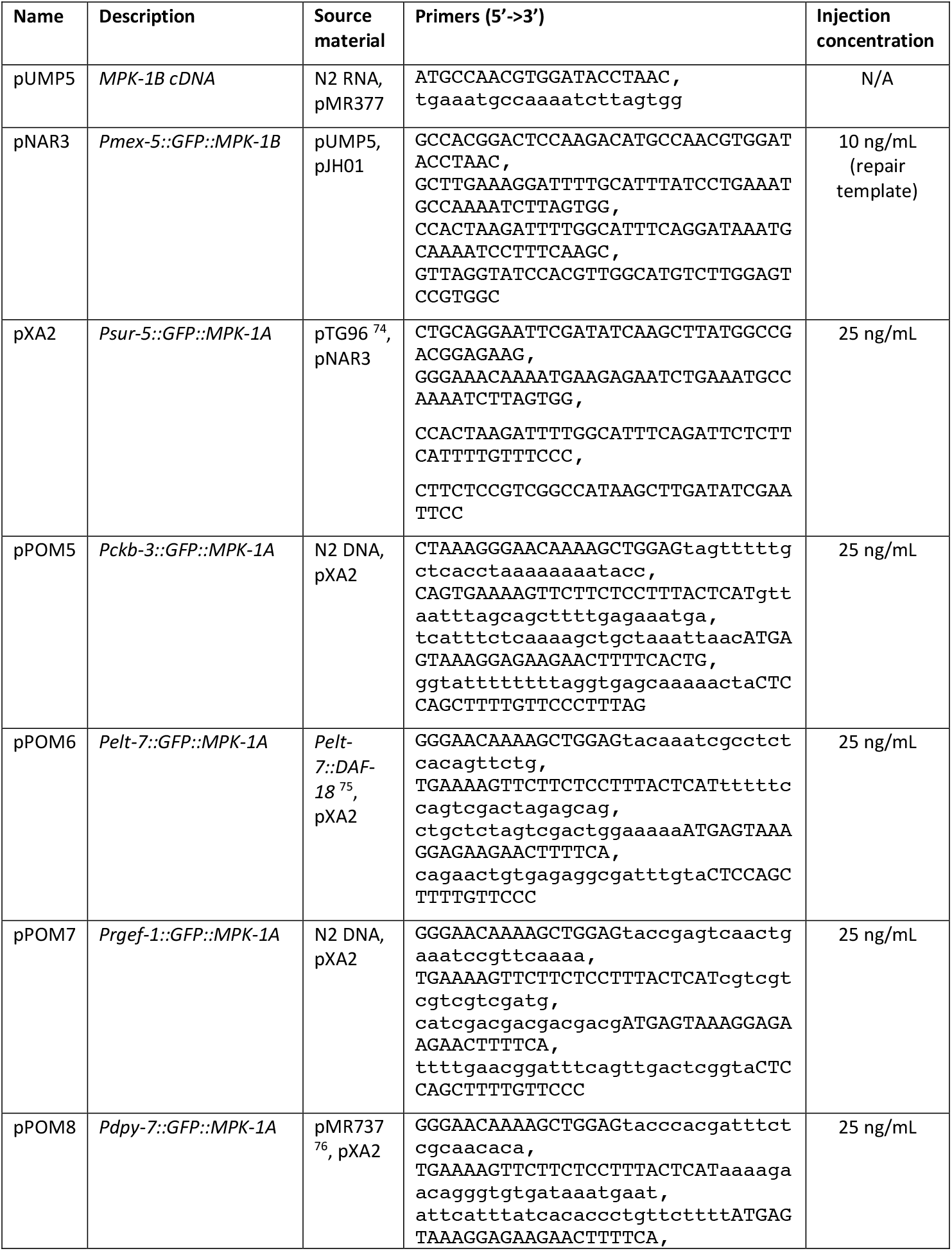
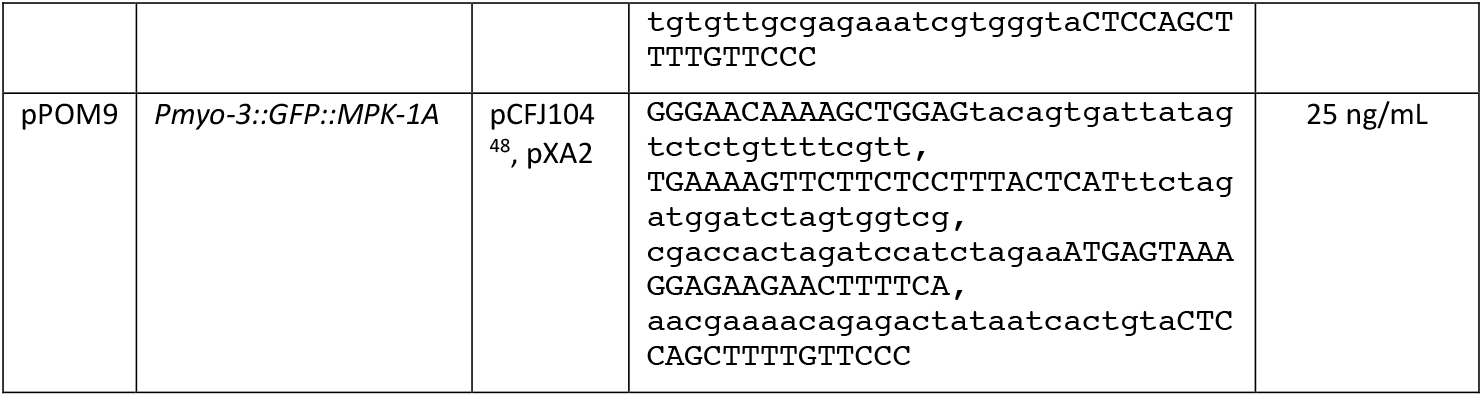
Plasmid design.

**Table S6.**
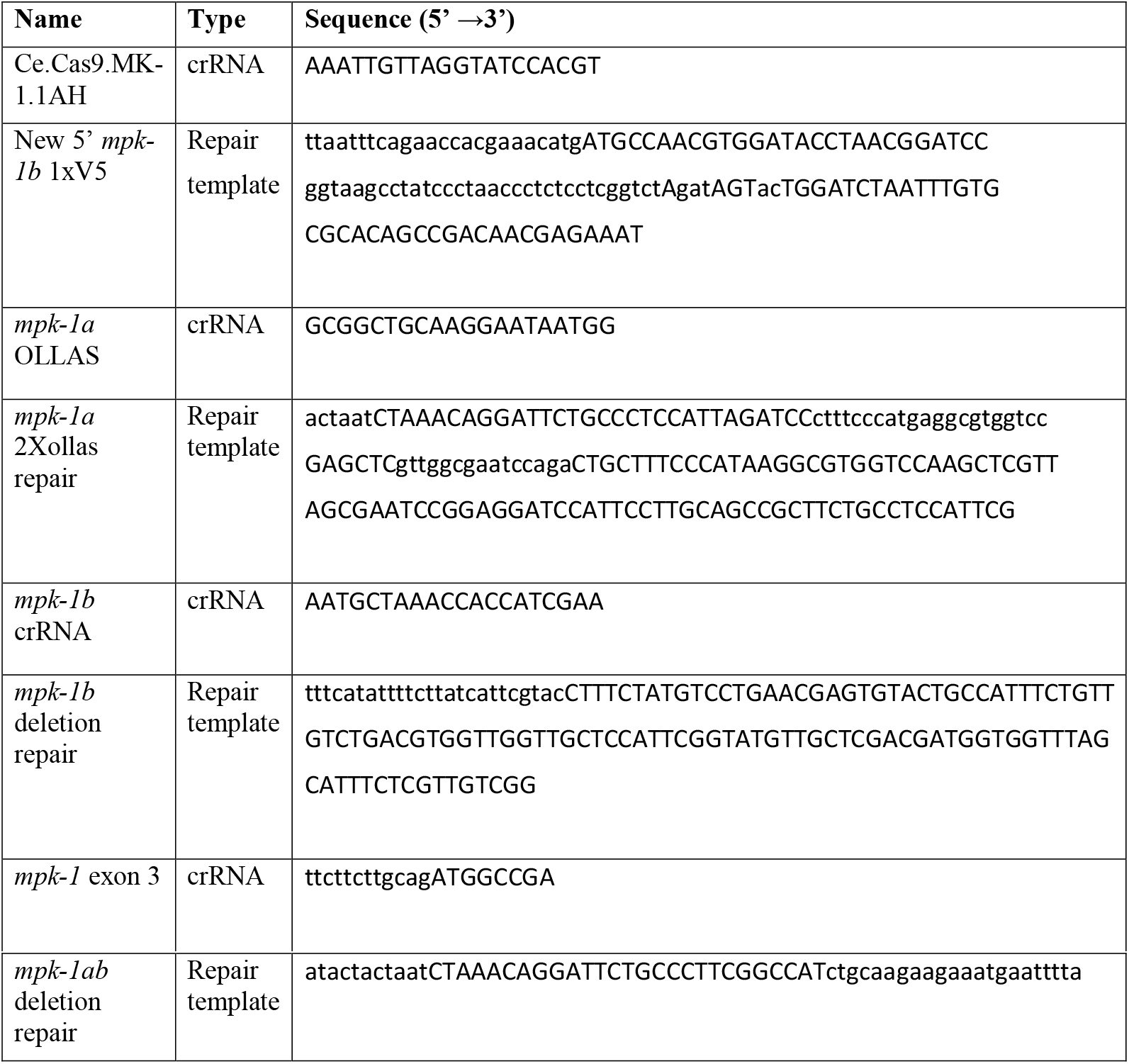
crRNA and repair sequences.

