## Supplementary figures and images for "Non-autonomous regulation of germline stem cell proliferation by somatic MPK-1/MAPK activity in *C. elegans*"

### Figure S1

MPK-1AB paper
Figure S1

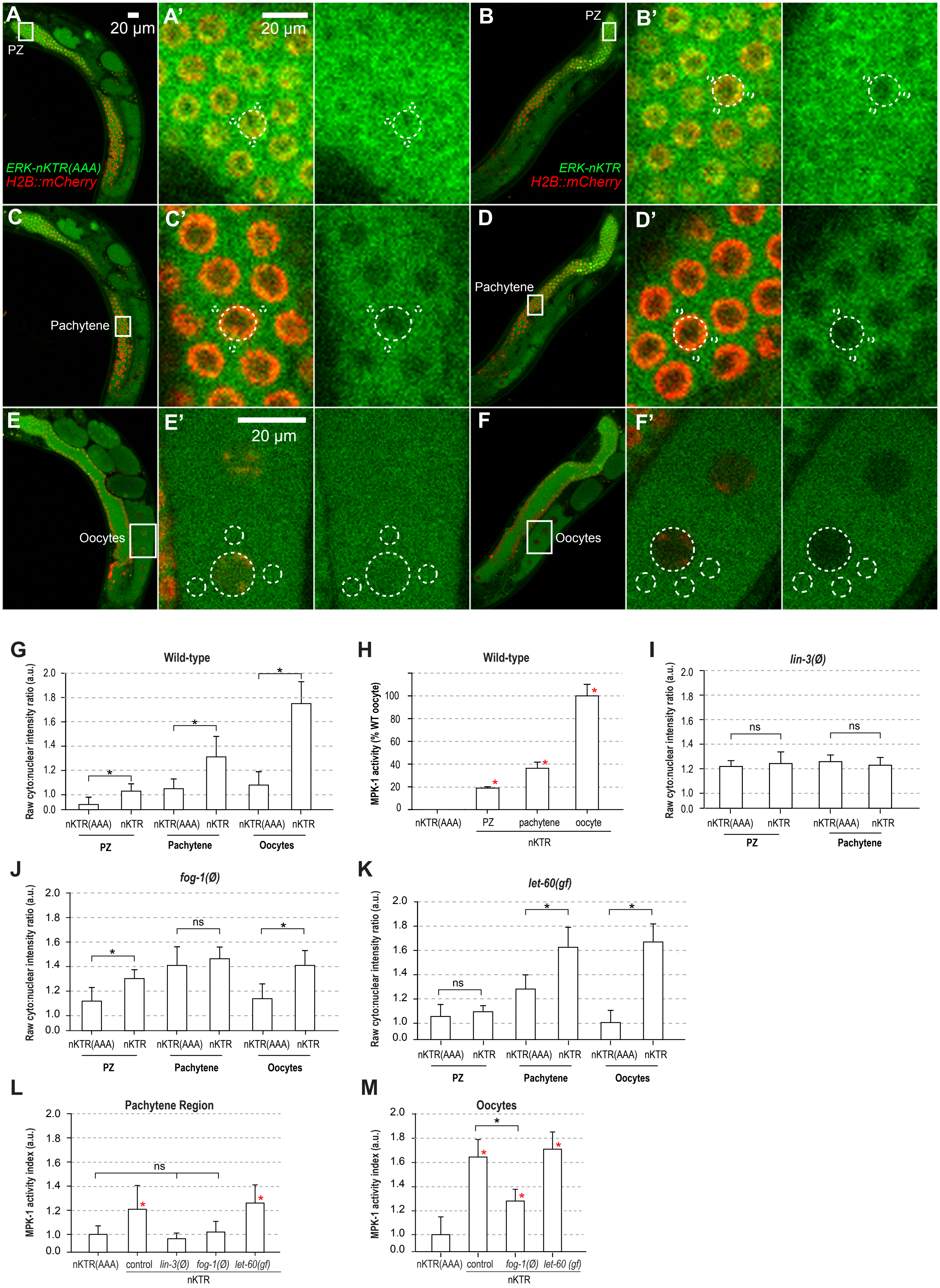

### Figure S2

MPK-1AB paper
Figure S2

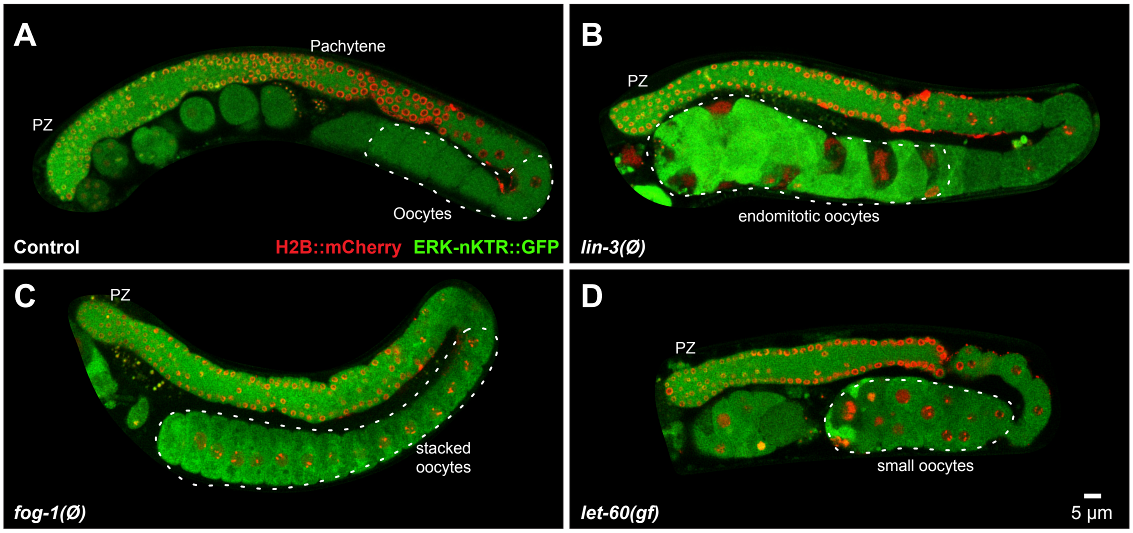

### Figure S3

MPK-1AB paper
Figure S3

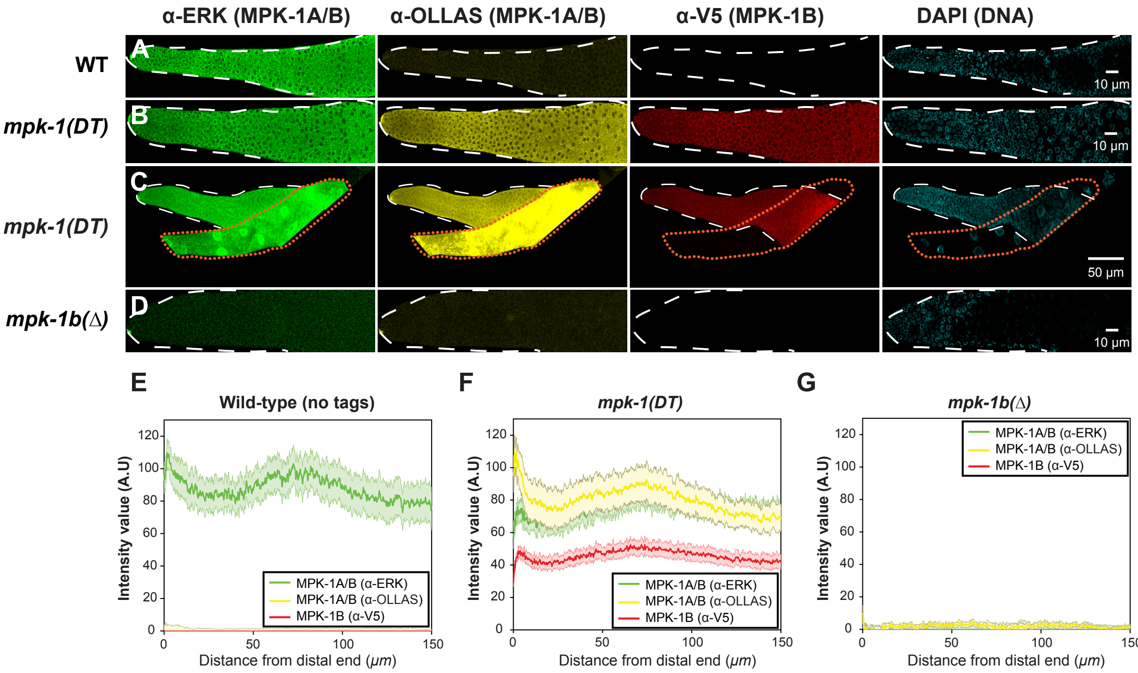


Lorem ipsum

### Figure S4

MPK-1AB paper
Figure S4

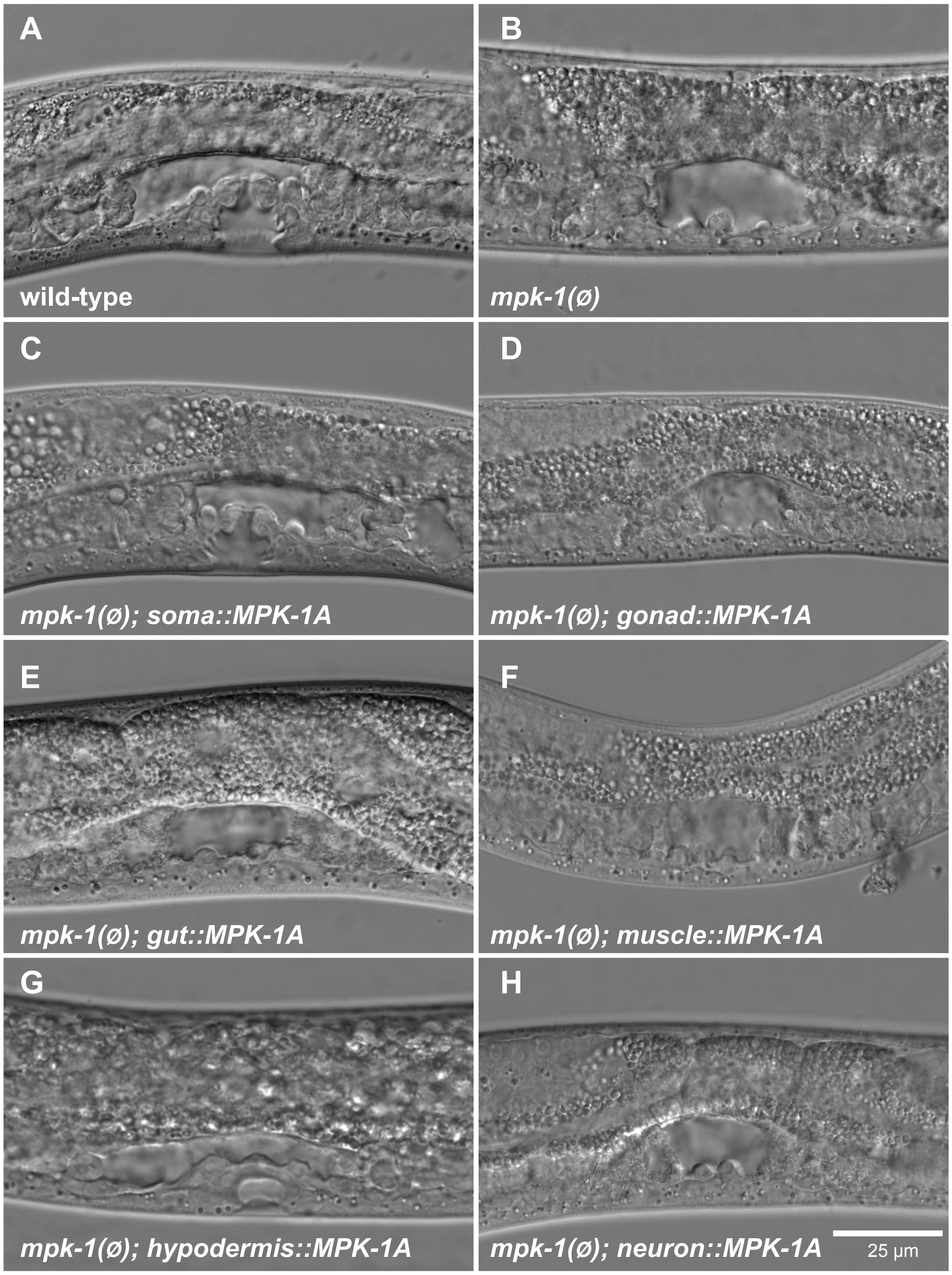
